# Hidden protein-altering variants influence diverse human phenotypes

**DOI:** 10.1101/2023.06.07.544066

**Authors:** Margaux L.A. Hujoel, Robert E. Handsaker, Maxwell A. Sherman, Nolan Kamitaki, Alison R. Barton, Ronen E. Mukamel, Chikashi Terao, Steven A. McCarroll, Po-Ru Loh

**Affiliations:** Division of Genetics, Department of Medicine, Brigham and Women’s Hospital and Harvard Medical School, Boston, MA, USA; Center for Data Sciences, Brigham and Women’s Hospital and Harvard Medical School, Boston, MA, USA; Program in Medical and Population Genetics, Broad Institute of MIT and Harvard, Cambridge, MA, USA; Stanley Center for Psychiatric Research, Broad Institute of MIT and Harvard University, Boston, MA, USA; Department of Genetics, Harvard Medical School, Boston, MA, USA; Computer Science and Artificial Intelligence Laboratory, Massachusetts Institute of Technology, Cambridge, MA, USA; Department of Biomedical Informatics, Harvard Medical School, Boston, MA, USA; Laboratory for Statistical and Translational Genetics, RIKEN Center for Integrative Medical Sciences, Yokohama, Japan; Clinical Research Center, Shizuoka General Hospital, Shizuoka, Japan; Department of Applied Genetics, School of Pharmaceutical Sciences, University of Shizuoka, Shizuoka, Japan

## Abstract

Structural variants (SVs) comprise the largest genetic variants, altering from 50 base pairs to megabases of DNA. However, SVs have not been effectively ascertained in most genetic association studies, leaving a key gap in our understanding of human complex trait genetics. We ascertained protein-altering SVs from UK Biobank whole-exome sequencing data (*n*=468,570) using haplotype-informed methods capable of detecting sub-exonic SVs and variation within segmental duplications. Incorporating SVs into analyses of rare variants predicted to cause gene loss-of-function (pLoF) identified 100 associations of pLoF variants with 41 quantitative traits. A low-frequency partial deletion of *RGL3* exon 6 appeared to confer one of the strongest protective effects of gene LoF on hypertension risk (OR = 0.86 [0.82–0.90]). Protein-coding variation in rapidly-evolving gene families within segmental duplications—previously invisible to most analysis methods—appeared to generate some of the human genome’s largest contributions to variation in type 2 diabetes risk, chronotype, and blood cell traits. These results illustrate the potential for new genetic insights from genomic variation that has escaped large-scale analysis to date.

## Introduction

Genomic structural variants (SVs), which modify 50 base pairs to megabases of DNA, account for the majority of base pairs of variation in each human genome^1^. Recent major efforts to study structural variation in human genomes have elucidated the landscape and mutational origins of SVs by ascertaining SVs from short-read sequencing of many thousands of individuals^2, 3^ and long-read sequencing of tens of individuals^4, 5^.

Assessing the impact of structural variation on human phenotypes requires genotyping SVs in large well-phenotyped cohorts. This has been possible for larger copy-number variants (CNVs) detectable from the SNP-array and whole-exome sequencing data generated at scale by biobank projects and consortia^6–11^. However, the effects of kilobase-scale and smaller SVs—which comprise the majority of SVs^1, 5^ —have remained largely hidden, requiring analyses of whole-genome sequencing data sets^12, 13^. Such analyses have demonstrated important influences of SVs on gene expression^14, 15^ but have only recently begun reaching the scale necessary to detect associations with human phenotypes^16–19^.

We sought to leverage population genetic principles to address this challenge. Studies of CNVs classically focused on large, extremely rare CNVs that recurred *ab initio* in different individuals or families^20^; most such CNVs affected many genes, making it hard to discern the mechanism by which they affected phenotypes. In contrast, far more CNVs are inherited by many people from common ancestors; these CNVs, which are generally smaller but can have disabling effects on specific, individual genes (and thus interpretable, specific effects on human biology), have often gone undetected. Since such CNVs are inherited by descent from common ancestors, we hypothesized that the additional information provided by SNP haplotypes^9, 21^ could enable analyses of abundant exome sequencing data to detect even small copy-number-altering SVs within individual protein-coding genes—including genes within multi-copy and segmental duplication regions. We applied this approach to explore the impacts of protein-altering SVs upon the ∼500,000 research participants in UK Biobank (UKB)^22, 23^.

## Results

### Haplotype-informed detection of rare protein-altering CNVs in UK Biobank

We first sought to sensitively detect rare protein-altering CNVs of all sizes—including CNVs that affected single exons—from UKB exome sequencing data (*n*=468,570). To enable detection at this resolution, we utilized a computational approach that can integrate information across individuals who share extended SNP haplotypes (Fig. 1a). Because CNVs inherited by descent from common ancestors will tend to be inherited on a shared SNP haplotype, analyzing such individuals together increases detection sensitivity (Fig. 1a). We previously used this approach to detect CNVs from genotyping array intensity data (while retaining sensitivity to larger *de novo* CNVs)^9^; here, we adapted the approach to model exome-sequencing read counts using negative binomial distributions with sample-and region-specific parameters (Methods). Importantly, leveraging haplotype-sharing information enabled analysis at 100bp resolution, allowing detection of small CNVs that only partially overlap single exons (Fig. 1a).

**Figure 1:**
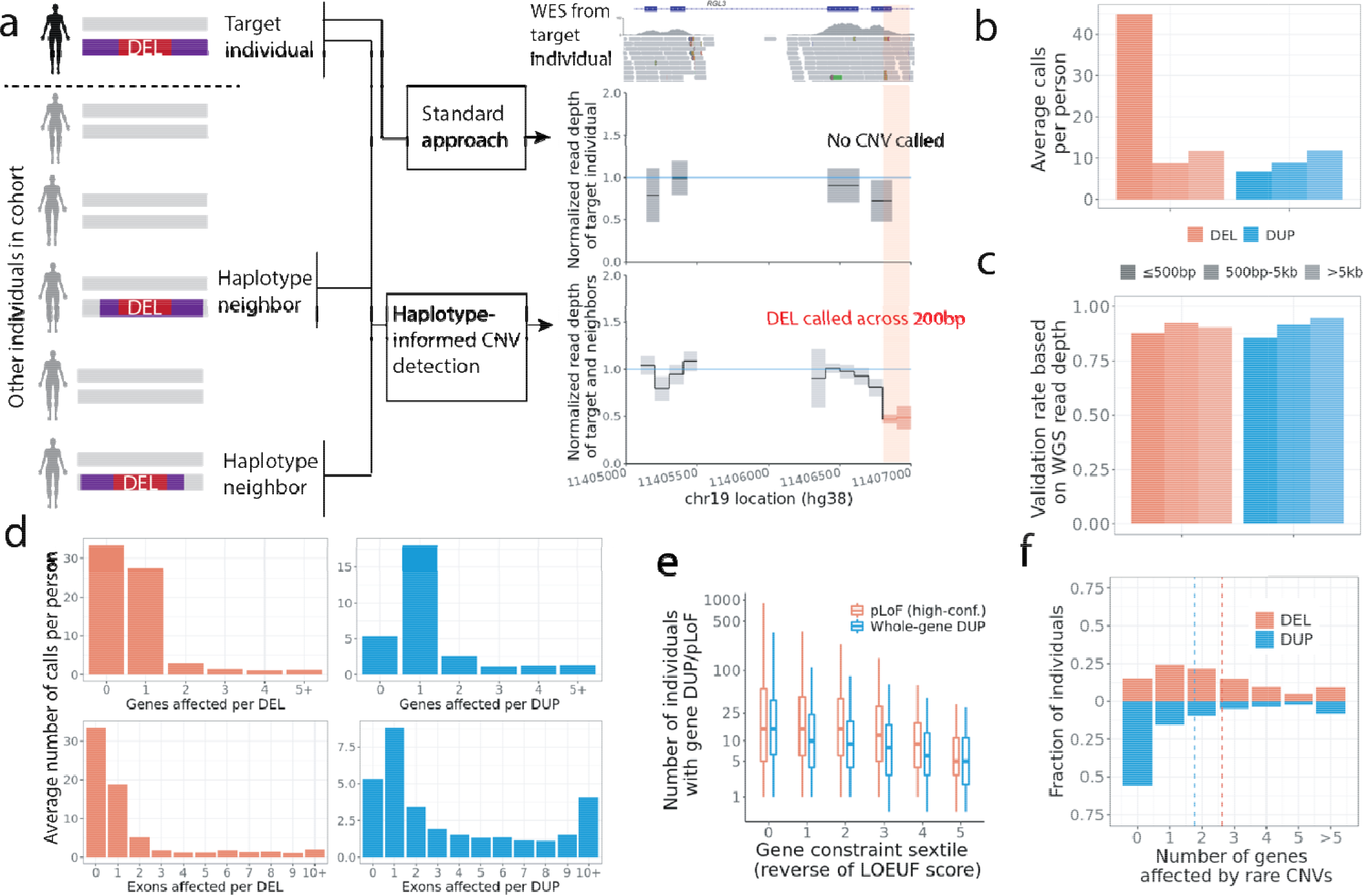
Haplotype-informed CNV detection from whole-exome sequencing in UK Biobank. (**a**) This approach improves power to detect CNVs by analyzing whole-exome sequencing read-depth data from an individual together with corresponding data from individuals sharing extended SNP-haplotypes (“haplotype neighbors”), facilitating analysis at the resolution of 100bp bins. In contrast, standard approaches analyze data from an individual alone, generally at exon-level resolution. (**b**) Average number of CNVs called per UKB participant, subdivided by copy-number change (deletion/duplication) and call length. (**c**) Validation rate of CNV calls based on analysis of whole-genome sequencing data for 100 UKB participants. (**d**) Average numbers of CNVs called per UKB participant affecting given numbers of genes or exons. (**e**) Distributions (across increasingly constrained gene sets) of observed counts of predicted loss-of-function deletions and whole-gene duplications in 487,205 UKB participants. Centers, medians; box edges, 25th and 75th percentiles; whiskers, 5th and 95th percentiles. (**f**) Fractions of UKB participants with given numbers of genes affected by rare CNVs.

We applied this approach to identify CNVs in the full UK Biobank cohort, focusing our main analyses on 454,682 European-ancestry participants to avoid confounding in subsequent association analyses. We identified an average of 93.4 CNVs per person (65.7 deletions and 27.7 duplications), roughly half of which were short deletions called across intervals of 500bp or less (Fig. 1b and Supplementary Table 1). This represented a twofold increase compared to a recent analysis of an interim UKB WES release (*n*=200K)^10^. Validation using whole-genome sequencing data for 100 participants indicated that false-positives were well controlled at <10%, with precision improving modestly with CNV size (Fig. 1c and Supplementary Table 2; Methods). Most deletions and roughly half of the duplications affected at most one exon (Fig. 1d), including some CNVs identified using only off-target reads that did not intersect any exons.

The most impactful variants were uncommon: across 18,651 genes, whole-gene duplications and CNVs predicted to cause loss-of-function (pLoF) were identified in a median of 8 and 11 individuals, respectively, with observed counts decreasing with increasing gene constraint (Fig. 1e). When focusing on genes rarely altered by such events, a mean of 4.4 genes per individual were affected (1.8 genes by whole-gene duplications and 2.6 genes by pLoF CNVs) (Fig. 1f), indicating improved sensitivity compared to state-of-the-art methods for rare CNV detection^24^.

### Rare large-effect CNVs implicate new gene-trait relationships

This resource of protein-altering copy-number variation in UK Biobank made it possible to discover new links between genetic variation and human phenotypes. To do so, we analyzed CNVs for association with 57 heritable quantitative traits (reflecting a broad spectrum of biological processes; Supplementary Data 1) using linear mixed models^25, 26^. We performed two sets of association analyses (Supplementary Fig. 1): (i) CNV-only analyses, which identified 180 CNV-trait associations (*P* < 5 x 10^−8^) likely to be driven by non-syndromic CNVs (Fig. 2a and Supplementary Data 2); and (ii) gene-level burden analyses that collapsed all types of pLoF variants (nonsyndromic CNVs, SNVs, and indels) to maximize power to detect rare loss-of-function effects. The burden analyses identified 100 pLoF gene-trait associations (*P* < 5 x 10^−8^) undetectable from analyses of pLoF SNVs and indels alone, demonstrating the benefit of incorporating CNVs in burden analyses (+20% increase in associations; Fig. 2b and Supplementary Data 3).

**Figure 2:**
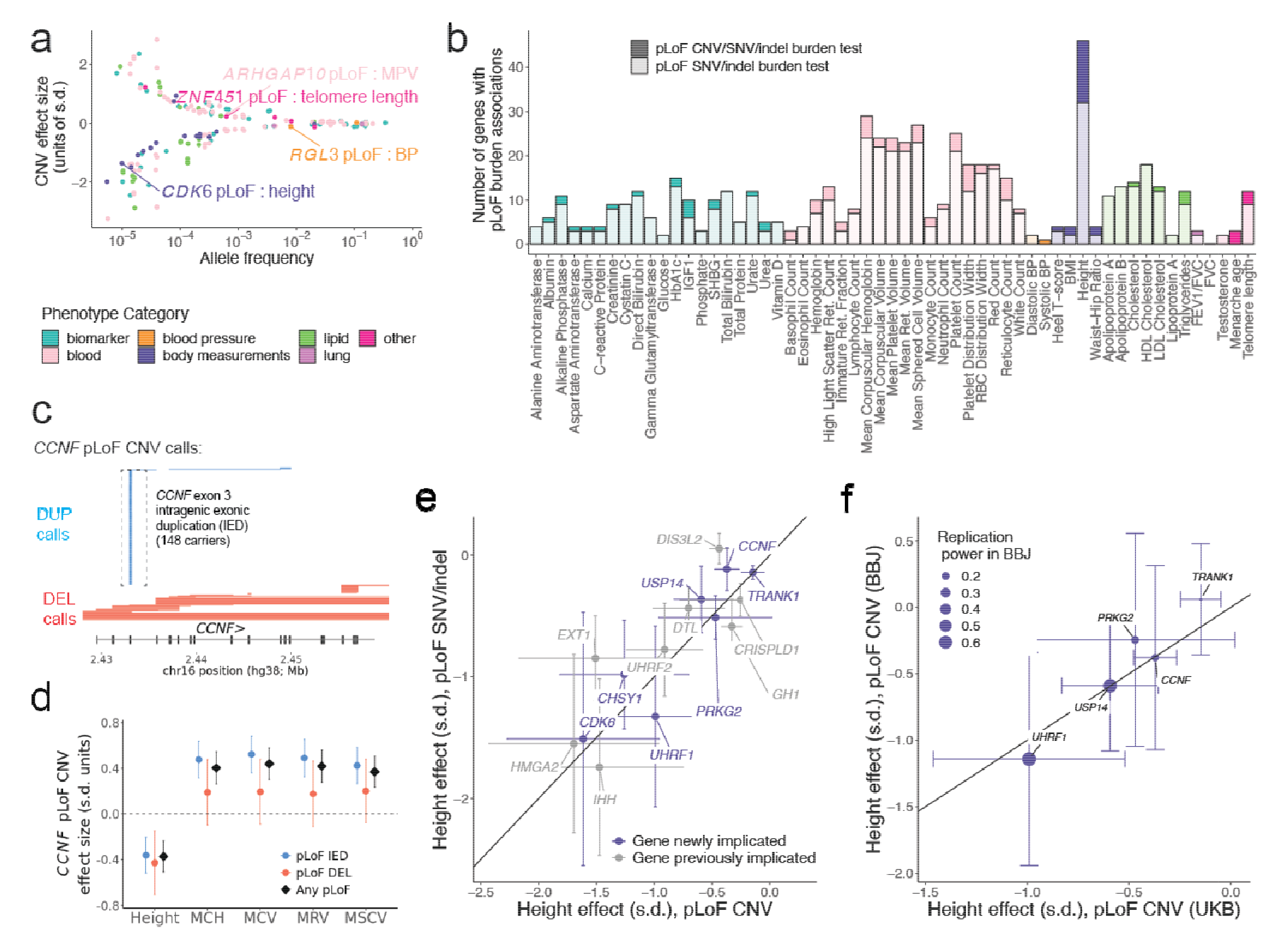
Association and fine-mapping analyses implicate rare large-effect CNVs and uncover new gene-trait relationships. (**a**) Effect size versus minor allele frequency for 180 likely-causal CNV-phenotype associations, colored by phenotype category. (**b**) Number of genes with pLoF burden associations (*P* < 5 x 10^-8^) per trait, colored by phenotype category, with darker shading corresponding to associations detectable only upon including pLoF CNVs (i.e., *P* > 5 x 10^-8^ for burden masks considering only SNVs and indels). (**c**) Genomic locations of *CCNF* pLoF CNV calls; boxed calls correspond to the rare duplication spanning a single 107bp exon. (**d**) Effect sizes of *CCNF* pLoF CNVs for height and erythrocyte traits. (**e**) Consistency of height effect sizes of pLoF CNVs with those of pLoF SNVs and indels. (**f**) Replication of height effect sizes of pLoF CNVs in BioBank Japan (for newly-implicated genes with at least five pLoF CNV carriers in BBJ). Error bars, 95% CIs.

Several of these associations implicated new gene-trait relationships, even for well-studied traits such as height for which common-variant association studies have reached saturation^27^. These included strong height-reducing effects (>1 s.d.) of ultra-rare pLoF variants (combined AF<0.0001) in *CHSY1*, which encodes an enzyme that synthesizes chondroitin sulfate (a structural component of cartilage), *UHRF1*, which encodes an E3 ubiquitin ligase that shares structural homology with UHRF2 (recently implicated by our previous work^9^), and *CDK6*, which harbors one of the strongest common-variant associations with height^28^. Rare pLoF variants in two other genes exhibited moderate height-reducing effects (–0.5 s.d.): *USP14*, which encodes a ubiquitin-specific protease, and *PRKG2*, which was recently implicated in autosomal recessive acromesomelic dysplasia^29^.

Another height association only discovered using CNVs involved *CCNF*, at which a rare duplication spanning a single 107bp exon accounted for more pLoF events than all other CNVs, SNVs, and indels combined (Fig. 2c). Validation using available UKB WGS data (*n*=200K) confirmed this CNV as a tandem duplication that was called from WES with 100% precision and 95% recall, illustrating the efficacy of haplotype-informed CNV detection (Supplementary Fig. 2a,1b and Supplementary Note). *CCNF* pLoF CNVs associated with a moderate decrease in height (–0.4 ± 0.1 s.d., *P* = 5.2 x 10^−12^) and appeared to have a pleiotropic effect on erythrocyte traits (Fig. 2d), motivating further study of this gene and its product, cyclin F.

While further work will be needed to confirm these findings and establish causality, two additional analyses provided evidence supporting their robustness. First, across the 15 height associations discovered only upon considering pLoF CNVs, the effect sizes of pLoF CNVs exhibited broad consistency with those of pLoF SNVs and indels (Fig. 2e), and this consistency held across traits (Supplementary Fig. 3). Second, for seven height-associated pLoF CNVs that affected genes not previously identified either by large-scale pLoF SNV/indel burden analyses^23, 30^ or CNV analyses^9^, we attempted replication in BioBank Japan^31^, observing broadly consistent effect sizes for the five genes with at least five pLoF CNV carriers in BBJ (Fig. 2f).

### *RGL3* loss of function associates with reduced hypertension risk

A low-frequency (AF=0.9%) deletion of part of exon 6 of the *RGL3* (Ral Guanine Nucleotide Dissociation Stimulator Like 3) gene associated with lower blood pressure (–0.11 ± 0.01 s.d.; *P* = 6.1 x 10^−23^) and decreased hypertension risk (OR = 0.86 [0.82–0.90]; Fig. 3a,b and Supplementary Table 3) as well as decreased serum calcium (−0.08 ± 0.01 s.d.; *P* = 6.0 x 10^-11^; Supplementary Data 2). Closer examination of this CNV showed it to be a 1.1kb deletion present in 8,117 UKB participants that intersects only 55bp of coding sequence (Fig. 3c and Supplementary Fig. 2c), yet had been successfully called with 99.9% precision and 88% recall (based on breakpoint-based follow-up analysis; Supplementary Note). This association replicated in the *All of Us* cohort (*n*=245,394) with a consistent decrease in hypertension risk (OR = 0.83 [0.75–0.92], *P* = 0.00026; Supplementary Fig. 2d and Supplementary Table 4a). The strongest blood pressure association at this locus was attained by a common *RGL3* missense variant (rs167479; AF= 47%) independent of the deletion (*R*^2^=0.005; Fig. 3a). Conditioning on rs167479 resulted in the deletion becoming the lead variant (Fig. 3a), supporting causality of both *RGL3* coding variants and explaining a previously-reported association of an intronic SNP in *RAB3D* (rs55670943, 76kb downstream^32^) that best tags the deletion (*R*^2^=0.66).

**Figure 3:**
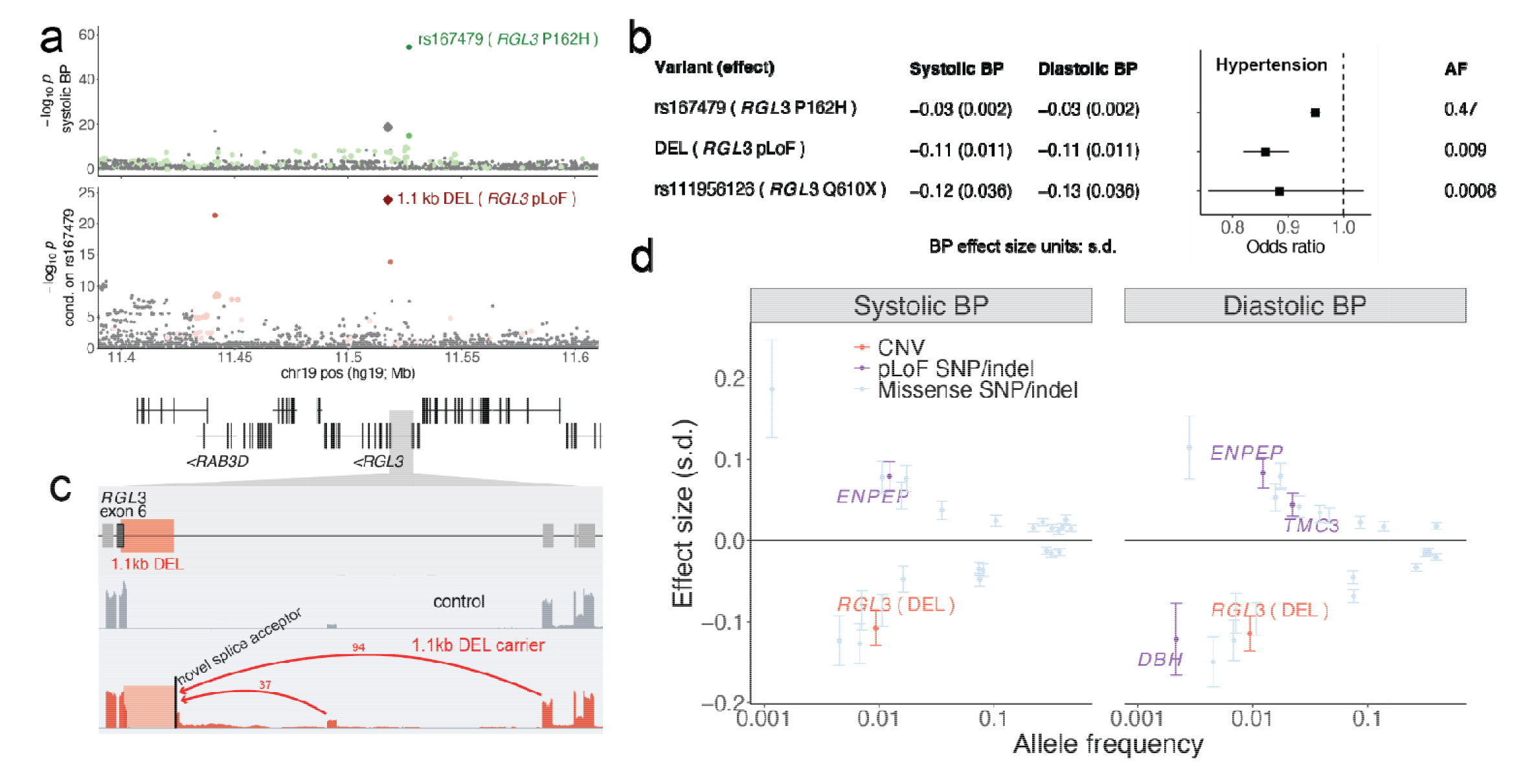
A low-frequency deletion in *RGL3* associates with reduced hypertension risk and generates novel splicing. (**a**) Associations of variants at the *RGL3* locus with systolic blood pressure in two steps of stepwise conditional analysis. Colored dots are variants in partial LD (*R*^2^ ≥ 0.01) with labeled variants. (**b**) Effect sizes and allele frequencies of a common *RGL3* missense variant (rs167479), the low-frequency 1.1 kb deletion, and a rare *RGL3* stop gain. (**c**) Evidence of novel *RGL3* splicing produced by the 1.1kb deletion. RNA sequencing read-depth data from GTEx are shown for a carrier of the deletion and a control sample (both thyroid); red arcs indicate novel splice junctions, labeled with counts of supporting RNA-seq reads. (**d**) Systolic and diastolic blood pressure effect sizes versus minor allele frequencies for nonsynonymous SNP and indel variants and the 1.1kb deletion. Error bars, 95% CIs.

The deletion variant appeared to have a much larger effect on blood pressure than the missense variant, similar to the effect of a rare *RGL3* stop gain (Fig. 3b), suggesting that it causes loss of RGL3 function. Analysis of RNA-sequencing data from the Genotype-Tissue Expression (GTEx) project^33^ provided insight into the transcriptional basis for this effect: carriers of the deletion, which removes the exon 6 splice acceptor, exhibited splicing into a novel splice acceptor upstream of the deletion (Fig. 3c), translating to an inframe substitution of a novel 23 amino acid sequence for a 19 amino acid segment of RGL3. Further work will be required to determine whether the modified protein is completely dysfunctional or the apparent LoF effect is mediated in part by reduced expression of *RGL3* alleles carrying the deletion (Supplementary Note).

Intriguingly, the blood pressure-lowering effect of the deletion in *RGL3* appears to be one of the strongest such effects among all coding variants genome-wide (Fig. 3d), and knockout of *RGL3* appears likely to be well-tolerated based on the presence of 37 UKB participants homozygous for the deletion who appeared to be generally healthy (Supplementary Note). These observations raise the possibility that RGL3, or a pathway in which it functions, could be an appealing target for antihypertensive drug development, motivating further study of RGL3 function.

### Identifying impacts of common coding copy-number polymorphisms

In addition to the genetic effects above, in uniquely mappable regions of the human genome, potentially important effects on human biology could arise within rapidly evolving gene families shaped by extensive recent gene duplication and divergence. The analytical technique above was designed to detect rare protein-altering CNVs within mappable regions. To enable exploration of common coding copy-number variation—including abundant variation within segmental duplications^34^—we developed another approach that first identifies genomic regions that harbor common copy-number-altering polymorphisms (based on correlated WES read-depth among parent-child trios) and then measures copy number in these regions by leveraging haplotype-sharing to denoise read-depth-derived estimates. This approach generalizes techniques we recently developed to study variable number tandem repeat (VNTR) polymorphisms^21^; here, we developed new algorithms to analyze a much larger set of CNV regions (Methods).

This approach detected 41,042 genomic regions (defined at the resolution of 100bp segments, exons, or previously-reported CNVs) with evidence of common copy-number-altering structural variation. These commonly copy-number-variable regions overlapped coding exons of 11% of autosomal genes, which tended to have lower probability of loss-of-function intolerance (average pLI=0.16 across such genes versus 0.25 across genes not impacted by common SVs; Supplementary Table 5).

Measuring copy-number variation in these regions—many of which are invisible to large-scale genetic analysis pipelines—provided a unique opportunity to search for associations with phenotypic variation in UK Biobank. Given the difficulty of modeling potentially-complex structural variation in such regions, compounded with the challenge of analyzing short-read alignments in low-mappability regions, we performed association analyses on quantitative, dosage-like measurements derived from read-depth rather than attempting to call discrete genotypes (Supplementary Fig. 1). We reasoned that while these measurements might only roughly represent structural variant alleles, association signals could still point to phenotypically-important SV regions meriting more careful follow-up.

This strategy proved fruitful: association analyses with 57 quantitative traits identified 375 associations at 99 loci not explainable by LD to nearby SNPs (Supplementary Data 4), recovering strong VNTR-phenotype associations we recently reported (including a 39bp coding repeat in *GP1BA* associated with platelet traits; *P* = 1.1 x 10^-133^ [ref. ^35^]), and revealing several new loci involving multi-copy variation poorly tagged by SNPs. Follow-up analyses of the most intriguing associations, detailed below, enabled further exploration of genetic variation at these loci and its influences on human health.

### Common coding variants hidden in segmental duplications modulate type 2 diabetes risk

Copy-number variation at 7q22.1 and in *CTRB2* associated with HbA1c and type 2 diabetes (T2D), contributing two of the top 20 T2D-associated loci in UKB (Fig. 4a). The 7q22.1 locus, which also generated the human genome’s strongest association with chronotype (Fig. 4b), contains a 99kb segmental duplication that is among the largest, most polymorphic CNVs in the human genome^5^ and encompasses four protein-coding genes (Fig. 4c). While copy number of the segment (typically ranging from 2–14 copies per diploid genome) associated with T2D status (*P* = 2.4 x 10^-13^ in UKB; Fig. 4c,d; replication *P* = 2.0 x 10^-4^ in *All of Us*; Supplementary Table 4b), we wondered whether this signal might be driven by paralogous sequence variants (PSVs): i.e., SNPs and indels carried on one or more copies of the 99kb segment within each allele. To genotype such variation, which is inaccessible to conventional analysis of short-read data, we first roughly estimated PSV genotypes from WGS read alignments (available for 200,018 UKB participants^18^) and then adapted our haplotype-informed approach to denoise PSV genotypes and impute them into the remainder of the UKB cohort (Supplementary Fig. 4; Methods).

**Figure 4:**
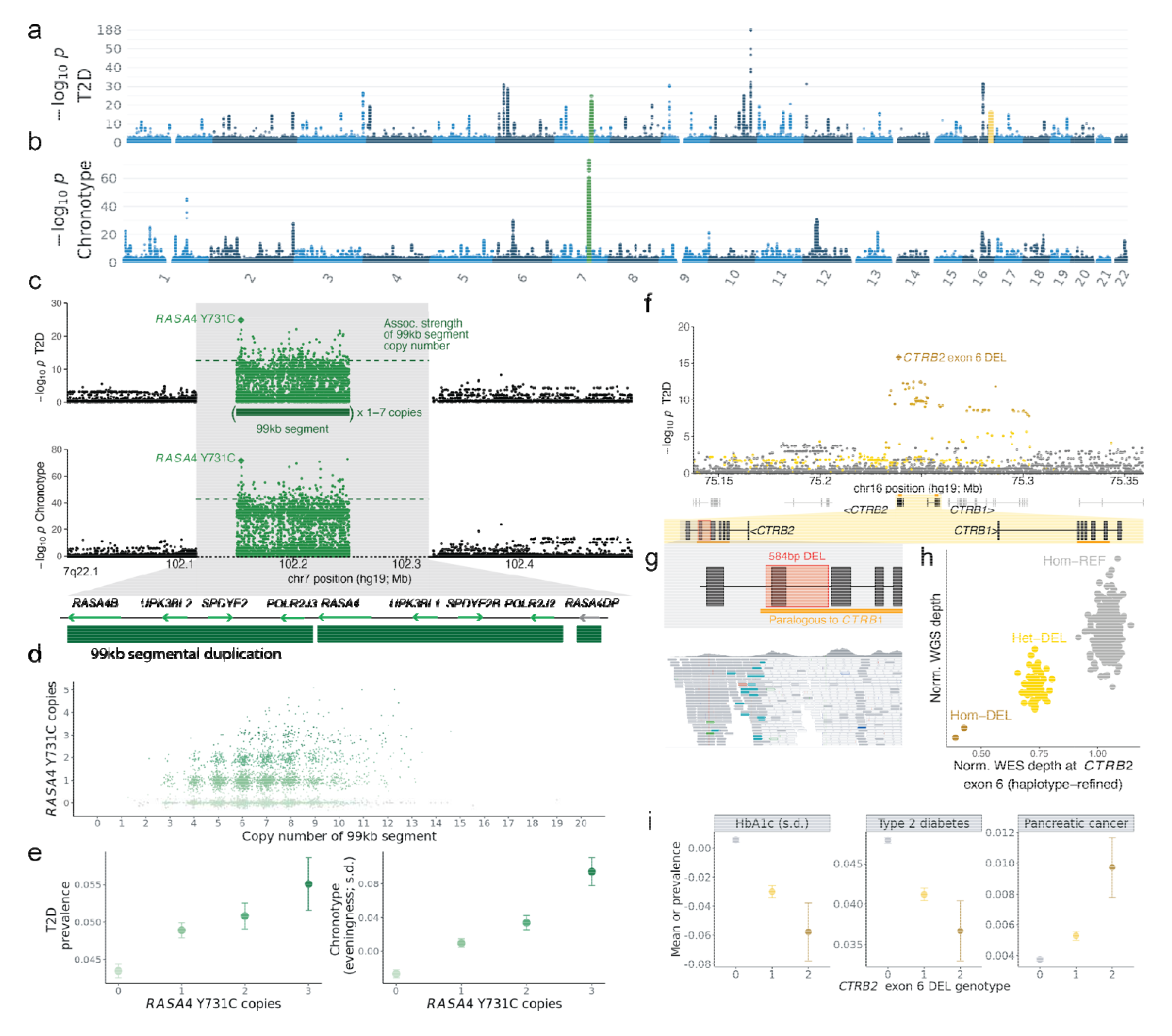
Coding variants within segmental duplications underlie top genetic associations with type 2 diabetes and chronotype. (**a**,**b**) Genome-wide associations with **(a)** type 2 diabetes (T2D) and **(b)** chronotype. (**c**) Associations of variation at 7q22.1 with chronotype and T2D. Associations of paralogous sequence variants (PSVs) within the 99kb repeat at this locus (1–7 copies per allele; 2 copies in GRCh37) are plotted in the center; green dashed line indicates association strength of copy number of the 99kb repeat. (**d**) Joint distribution of copy-number estimates for the 99kb segmental duplication and the *RASA4* Y731C missense variant. (**e**) T2D prevalence and mean chronotype (in standardized units; higher for “evening people”) as a function of number of copies of the *RASA4* Y731C missense varia t. (**f**) T2D associations at the *CTRB2* locus; colored dots are variants in partial LD (*R*^2^ > 0.01) with the *CTRB2* exon 6 deletion. (**g**) Location of the 584bp deletion spanning *CTRB2* exon 6 (top) and exome-sequencing read alignments for a deletion carrier (bottom); most reads aligned to the region paralogous to *CTRB1* do not map uniquely and are colored white. (**h**) Scatter plot of normalized whole-genome and whole-exome sequencing read depths at *CTRB2* exon 6. (**i**) Mean HbA1c and prevalence of T2D and pancreatic cancer as a function of *CTRB2* exon 6 deletion genotype. Error bars, 95% CIs.

Intriguingly, testing PSVs at 7q22.1 for association with T2D and chronotype identified a common missense PSV in *RASA4* (encoding Ras GTPase-activating protein 4) as the most strongly associated variant for T2D and second-strongest for chronotype (*P* = 1.3 x 10^-25^ and 2.6 x 10^-72^, respectively; Supplementary Table 6a; T2D replication *P* = 2.8 x 10^-5^ in *All of Us*; Supplementary Table 4b). For both phenotypes, the number of copies of *RASA4* with this mutation (encoding a Y731C substitution in the canonical transcript) associated much more strongly than copy number of the 99kb segment (Fig. 4c), and for chronotype, the *RASA4* missense PSV associated far more strongly than variants at all other loci across the genome (Fig. 4b). The contribution of this locus to each phenotype had largely been hidden from previous analyses, as SNPs flanking the segmental duplication poorly tag multi-copy variation within it (Fig. 4c). The total number of copies of *RASA4* carrying the Y731C missense PSV (typically ranging from 0 to 3 per individual; Fig. 4d) associated with increasing T2D risk and “eveningness” (i.e., later preferred bedtime/rising time) (Fig. 4e), with a 1.30-fold (1.21–1.39) range in odds of T2D. This PSV appears to be a strong candidate causal variant given its protein-altering effect and support from statistical fine-mapping (Supplementary Note); however, further study will be required to determine whether it indeed underlies one or both of these associations, and if so, how this mutation affects *RASA4* function.

The *CTRB2* gene encodes the chymotrypsinogen B2 protein, which is primarily produced in the pancreas, is converted into the active enzyme chymotrypsin B through enzymatic cleavage in the small intestine, and plays an important role in the digestive process^36^. A common 584bp deletion (AF=0.08) spanning exon 6 of *CTRB2* appeared to underlie another top locus for T2D (Fig. 4a,f). This deletion falls within a region of high homology to *CTRB1*, but our analysis pipeline successfully captured the copy-number variability of exon 6 from WES read-depth despite the low mappability (Fig. 4g,h). The deletion associated with decreased T2D risk (*P* = 1.6 x 10^-16^, strongest at the locus; OR = 0.86 [0.82–0.89]), replicating in *All of Us* (*P* = 2.3 x 10^-5^; Supplementary Fig. 2e and Supplementary Table 4c). We also replicated a recently-reported association of the deletion (which was shown to impair chymotrypsin B2 function and localization) with increased risk of pancreatic cancer^37^ (*P* = 4.2 x 10^-12^; Fig. 4i and Supplementary Table 6b). The opposite effect direction of these associations is notable given the overall epidemiological association of T2D with increased pancreatic cancer risk^38^.

### *FCGR3B* and *DEFA1A3* segmental duplication variants associate strongly with blood traits

Copy-number polymorphisms in two other segmental duplication regions produced two of the top five independent associations with count of basophils (Fig. 5a), a type of white blood cell that plays a role in the immune response and the regulation of allergic reactions. Here our analysis helped to recognize powerful effects within the *FCGR3* gene family, which encodes a family of cell surface receptors found on various immune cells, including neutrophils, macrophages, and natural killer cells; FCGRs play a crucial role in the immune response by recognizing and binding to the Fc portion of immunoglobulins (antibodies) that are bound to antigens^39^. In UKB, copy number of *FCGR3B* (which our analysis disambiguated from that of its paralog, *FCGR3A*) associated strongly with increased basophil count (*P* = 1.4 x 10^-82^, far exceeding the associations of nearby SNPs, which poorly tagged the recurrent CNV; Fig. 5b,c). Analysis of FCGR3B plasma protein levels corroborated the *FCGR3B* genotypes (Supplementary Fig. 5). *FCGR3B* deletion has previously been associated with several autoimmune disorders^40, 41^; here, decreasing *FCGR3B* gene dosage also associated with increasing risk of chronic obstructive pulmonary disease (*P* = 7.5 x 10^-7^; Fig. 5d and

**Figure 5:**
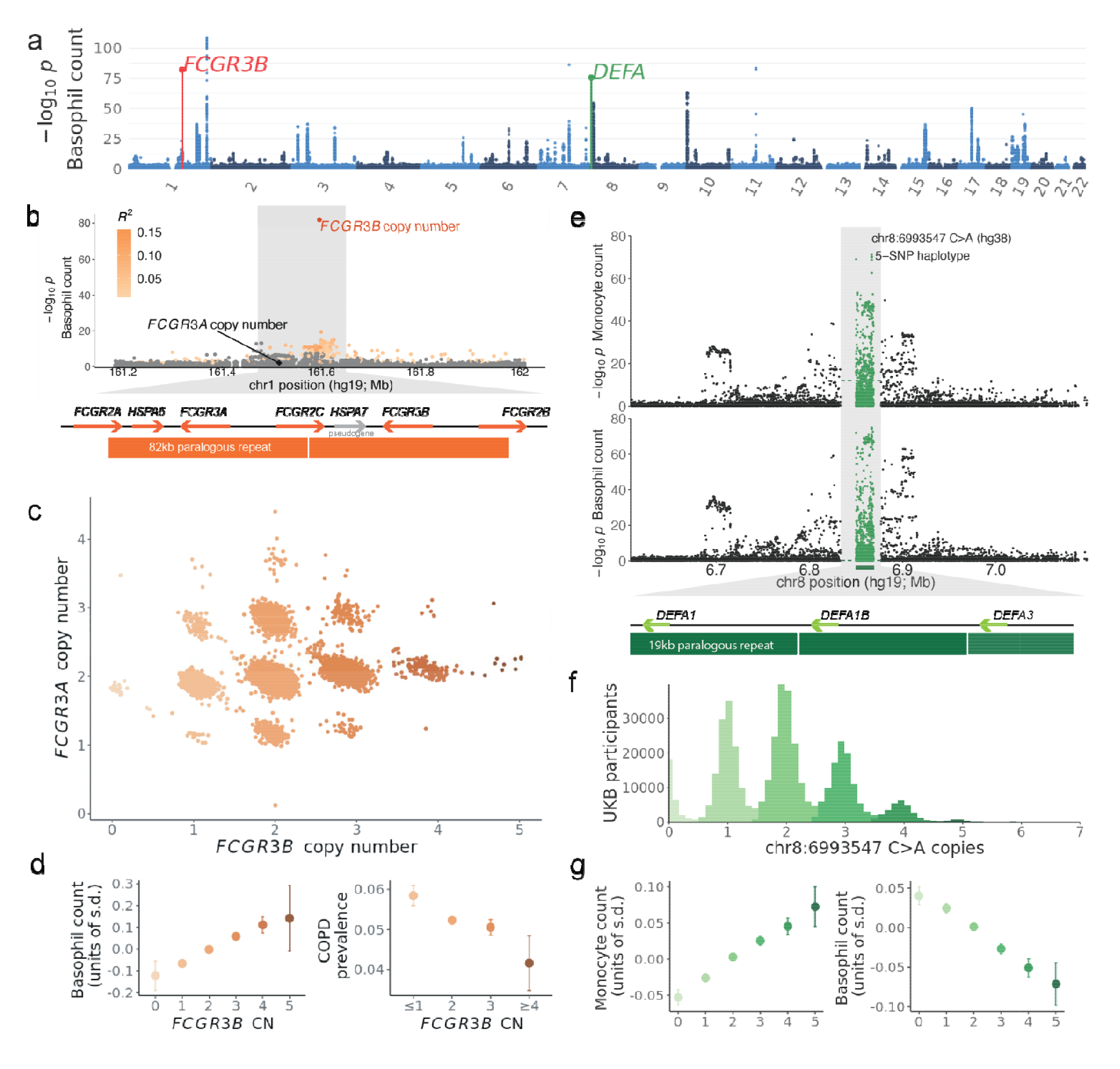
Variation in segmental duplications generates two of the top five genetic associations with basophil counts. (**a**) Genome-wide associations with basophil counts. (**b**) Associations with basophil counts at the *FCGR3B* locus; colored dots are variants in partial LD (*R*^2^ > 0.01) with *FCGR3B* copy number. (**c**) Joint distribution of copy-number estimates for *FCGR3A* and *FCGR3B*. (**d**) Mean basophil count and prevalence of chronic obstructive pulmonary disease (COPD) as a function of *FCGR3B* copy number. (**e**) Associations with basophil counts at the *DEFA1/DEFA3* locus. PSVs within the 19kb repeat at this locus are plotted as in Fig. 4c; green dashed line indicates association strength of copy number of the 19kb repeat. (**f**) Histogram of the number of copies of the 19kb repeat carrying the 5-SNP haplotype represented by chr8:6993547 C>A (GRCh38 coordinates). (**g**) Mean monocyte and basophil count as a function of copy number of the 5-SNP haplotype. Error bars, 95% CIs.

Supplementary Table 6c). The *FCGR* locus on 1q23.3 is known to contain many functional variants including multiple distinct CNVs^39^, such that while the basophil count association appears to be driven by *FCGR3B* copy number, other associations at this locus (Supplementary Data 4) may reflect other causal variants.

We were also able to recognize potent effects within the family of alpha-defensin genes, a rapidly evolving gene family that encodes a class of small, cationic peptides that are part of the innate immune system and play a crucial role in host defense against microbial infections^42^. Variation at the alpha-defensin gene cluster at 8p23.1 associated strongly with basophil count (Fig. 5a,e) as well as monocyte count (Fig. 5e). Alleles at this locus contain a highly variable number of copies of a 19kb repeat, each containing a single alpha-defensin gene^43^. Analysis of PSVs within this region (which had not previously been studied at scale, similar to the *RASA4* locus at 7q22.1) suggested that the number of copies of the 19kb segment carrying a tightly-linked 5-SNP haplotype within an Alu element inside the repeat—rather than the total number of copies of the repeat—might drive the association (Fig. 5e and Supplementary Table 6d). The number of copies of this repeat type typically ranged from 0 to 5 per individual (Fig. 5f) and associated with steadily increasing monocyte count and decreasing basophil count (Fig. 5g); however, we caution that a functional consequence of the 5-SNP haplotype is not immediately clear, unlike for the protein-coding variants at other loci we have highlighted.

### *SIGLEC14/5* gene fusion demonstrates tissue-specific promoter activity

A common, pleiotropic CNV at the *SIGLEC14–SIGLEC5* locus provided a unique opportunity to isolate a tissue-specific effect of a promoter element. A deletion allele at this locus that is particularly common in East Asians creates a fusion gene in which *SIGLEC5* is placed under the control of the *SIGLEC14* promoter^44^ (Fig. 6a,b). In UK Biobank, this CNV associated with several blood cell indices and serum biomarkers (*P* = 1.5 x 10^-8^ to 1.7 x 10^-37^; Fig. 6c; Supplementary Data 4 and Supplementary Table 7a). Follow-up analysis in GTEx revealed an unusually tissue-specific effect of the fusion on *SIGLEC5* expression, with the effect size varying greatly in magnitude and even direction across tissues (Fig. 6d). This phenomenon appeared to be explained by the further observation that the fusion’s tissue-specific effects on *SIGLEC5* expression tracked with relative efficiency of the *SIGLEC14* and *SIGLEC5* promoters across tissues (measured by the relative expression of *SIGLEC14* and *SIGLEC5* in individuals homozygous for the reference allele), such that the variable effect of the fusion was in fact consistent with its substitution of the *SIGLEC14* promoter in place of the *SIGLEC5* promoter (Fig. 6d and Supplementary Table 7b).

**Figure 6:**
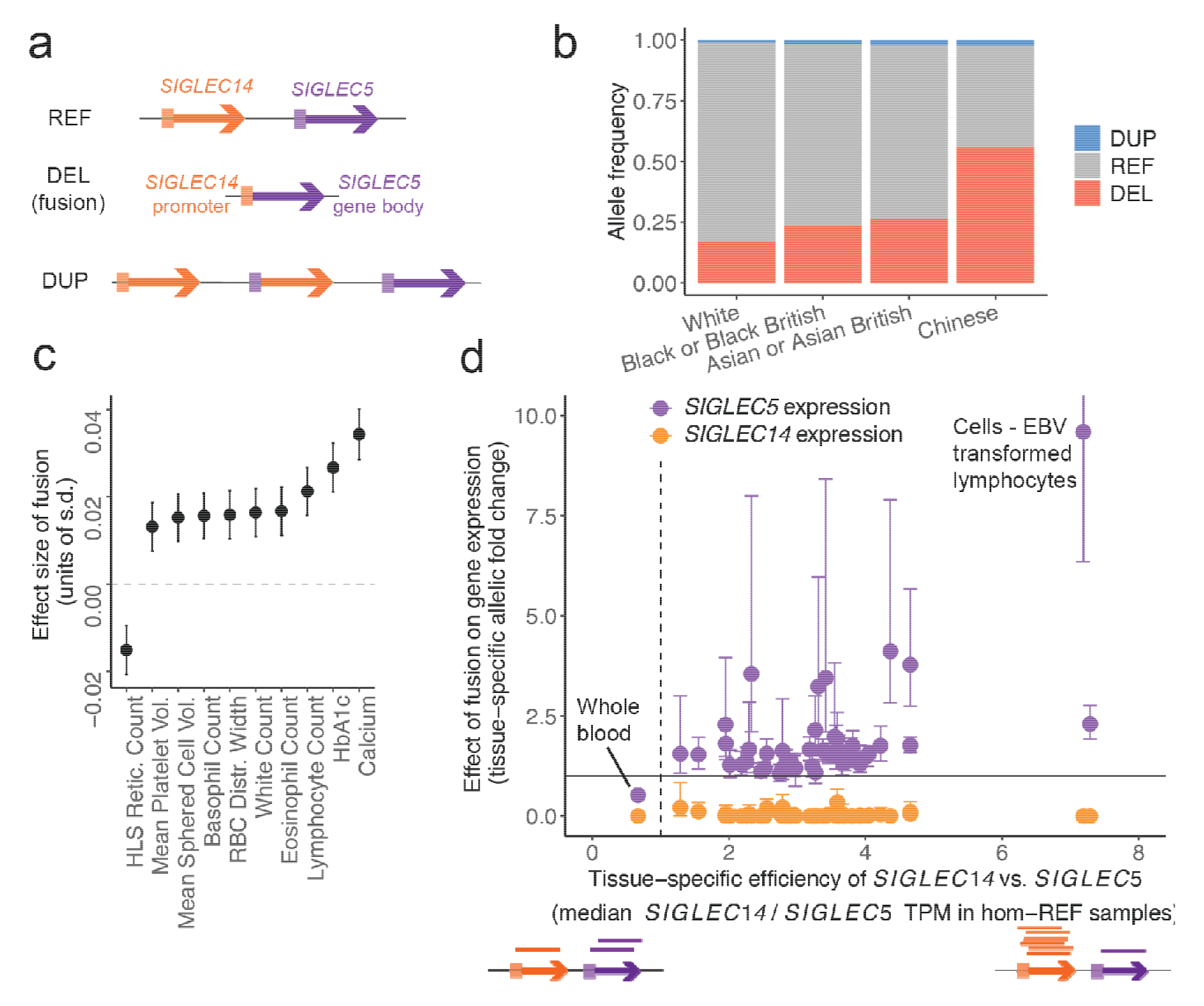
Common, pleiotropic *SIGLEC14–SIGLEC5* gene fusion illustrates tissue-specific promoter activity. (**a**) Gene diagram of *SIGLEC14* and *SIGLEC5*. A common deletion allele fuses the *SIGLEC14* promoter to the *SIGLEC5* gene body, and the reciprocal duplication allele is also observed at lower frequencies. (**b**) Allele frequency of gene fusion and duplication events in UK Biobank, stratified by reported ethnicity. (**c**) Effect size of fusion on blood indices and serum biomarker traits. (**d**) Allelic fold change effect of fusion on *SIGLEC5* and *SIGLEC14* gene expression across GTEx tissues tracks with relative efficiency of *SIGLEC14* promoter vs. *SIGLEC5* promoter in each tissue. Error bars, 95% CIs.

Other notable results included two strong associations with leukocyte telomere length, one involving an 84bp deletion within an alternative last exon of *ZNF208* (*P* = 1.7 x 10^-53^), and the other involving difficult-to-resolve copy number variation in the *CLEC18A/B/C* gene family (*P* = 1.0 x 10^-40^), which exhibits complex structural variation across two loci >4Mb apart^16^. Future analyses of long-read data sets will be better able to probe variation at such segmental duplications and elucidate phenotypic consequences hinted at here.

## Discussion

These results illustrate the phenotypic impact of protein-altering copy-number polymorphisms hidden from large-scale analyses to date. Here we observed that such variants include top genetic influences on human phenotypes that have eluded genetic association studies despite steadily increasing sample sizes and phenotyping precision. We further identified new gene-trait relationships implicated by rare CNVs that, for many genes, comprise a substantial proportion of loss-of-function events. We do caution that some of these associations still need replication; here we replicated a subset of the associations and observed corroborating evidence from allelic series for others. Additionally, while the protein-coding variants that we have implicated have clear effects on amino acid sequence or gene dosage, experimental work is needed to confirm causality of these variants and understand how they influence function and ultimately phenotype.

Our analyses here, based on exome sequencing of 468,570 individuals in UK Biobank, are far from comprehensive. While our haplotype-informed approach accurately recognized several sub-exonic CNVs that we linked to phenotypes, we expect that it missed very rare, short CNVs carried by only a few UKB participants. We also did not attempt to study shorter tandem repeats, which require specialized techniques^45^. Additionally, our analysis of common CNV regions—via rough quantifications of copy number—imperfectly modeled complex, multi-allelic structural variation. More precise genotyping of variation in such regions is needed, particularly in segmental duplications (∼7% of the human genome^46^). Our analyses were also limited in scope by the generally healthy, predominantly European-ancestry composition of the UK Biobank cohort. A search for associations between disease traits and gene-inactivating variants (including CNVs) only recovered known Mendelian disease genes (Supplementary Data 5), reflecting limited power to study rare diseases in population cohorts. Finally, while we prioritized compelling associations to highlight here using a stringent statistical fine-mapping filter, relaxing this filter would yield many more associations.

We anticipate that expanding genome-sequencing projects, including some that will use long reads^17, 47^, will overcome many of these limitations in the coming years, and we look forward to further insights into phenotypic consequences of both coding and noncoding structural polymorphisms in the years ahead.

## Acknowledgments

We thank Dr. Yukihide Momozawa in RIKEN Center for Integrative Medical Sciences and the members of the BioBank Japan Project, headquartered in the University of Tokyo Institute of Medical Science, for supporting this project. This research was conducted using the UK Biobank Resource under application number 40709. M.L.A.H. was supported by US NIH Fellowship F32 HL160061. R.E.H. and S.A.M. were supported by US NIH grant R01 HG006855. M.A.S. was supported by US NIH Fellowship F31 MH124393. N.K. was supported by US NIH training grant T32 HG002295. A.R.B. was supported by US NIH fellowship F31 HL154537. R.E.M. was supported by US NIH grant K25 HL150334. C.T. was supported by Japan Agency for Medical Research and Development (AMED) grants JP21kk0305013, JP21tm0424220, and JP21ck0106642, and Japan Society for the Promotion of Science (JSPS) KAKENHI grant JP20H00462. P.-R.L. was supported by US NIH grant DP2 ES030554, a Burroughs Wellcome Fund Career Award at the Scientific Interfaces, and the Next Generation Fund at the Broad Institute of MIT and Harvard. The funders had no role in study design, data collection and analysis, and decision to publish or preparation of the manuscript. Computational analyses were performed on the O2 High Performance Compute Cluster, supported by the Research Computing Group, at Harvard Medical School (http://rc.hms.harvard.edu), the UK Biobank Research Analysis Platform, and the *All of Us* Researcher Workbench. The *All of Us* Research Program is supported by the National Institutes of Health, Office of the Director: Regional Medical Centers: 1 OT2 OD026549; 1 OT2 OD026554; 1 OT2 OD026557; 1 OT2 OD026556; 1 OT2 OD026550; 1 OT2 OD 026552; 1 OT2 OD026553; 1 OT2 OD026548; 1 OT2 OD026551; 1 OT2 OD026555; IAA #: AOD 16037; Federally Qualified Health Centers: HHSN 263201600085U; Data and Research Center: 5 U2C OD023196; Biobank: 1 U24 OD023121; The Participant Center: U24 OD023176; Participant Technology Systems Center: 1 U24 OD023163; Communications and Engagement: 3 OT2 OD023205; 3 OT2 OD023206; and Community Partners: 1 OT2 OD025277; 3 OT2 OD025315; 1 OT2 OD025337; 1 OT2 OD025276. In addition, the *All of Us* Research Program would not be possible without the partnership of its participants. The Genotype-Tissue Expression (GTEx) Project was supported by the Common Fund of the Office of the Director of the National Institutes of Health, and by NCI, NHGRI, NHLBI, NIDA, NIMH, and NINDS.

## Declaration of Interests

The authors declare no competing interests.

## Methods

### UK Biobank genetic data

Whole-exome sequencing (WES) data was previously generated for ∼470,000 UK Biobank participants^23^. We analyzed these data together with SNP-haplotypes we previously generated for 487,409 participants^48^ in the UKB SNP-array and imputation data set (imp_v3)^22^. We performed haplotype-informed CNV detection on all UK Biobank participants with SNP haplotypes (including 468,570 participants with WES data passing QC as well as the remaining ∼3% of the imp_v3 samples via an imputation approach; Supplementary Note). We also analyzed whole-genome sequencing (WGS) data available for 200,018 participants^18^ in validation analyses and follow-up analyses of paralogous sequence variants within segmental duplications. We focused our primary analyses on individuals of self-reported White ethnicity, excluding individuals with trisomy 21, blood cancer, aberrantly many CNV calls, and those who had withdrawn at the time of our study (Supplementary Note), resulting in 454,682 participants being included in main analyses.

### UK Biobank phenotype data

We primarily analyzed 57 heritable quantitative traits measured on most UK Biobank participants (Supplementary Data 1), including 56 quantitative traits we recently analyzed^9^ along with telomere length. We reprocessed blood traits using a slightly modified pipeline in which we did not perform outlier removal (because some rare variants produce extreme blood indices): i.e., within strata of sex and menopause status, we performed inverse normal transformation and then regressed out age, ethnicity, alcohol use, smoking status, height, and BMI^9^. We processed the telomere length phenotype (Data-Field 22192)^49^ by applying inverse normal transformation. The remaining traits were processed as previously described^9^.

In secondary analyses (e.g., follow-up at loci of interest), we analyzed additional traits including binary disease outcomes derived from self-report (touchscreen questionnaire at assessment), hospital inpatient records, and cancer and death registries as well as plasma protein abundances for FCGR3B. In particular, we analyzed hypertension (174,773 cases and 279,891 controls; first-occurrence Data-Field 131286), type 2 diabetes (21,292 cases and 432,324 controls; derived from self-reported doctor-diagnosed T2D Data-Field 2443, following ref.^50^), pancreatic cancer (1,816 cases and 452,848 controls; ICD-10 code C25 from hospital records and cancer and death registries), and COPD (23,875 cases and 430,789 controls; first-occurrence Data-Field 131492). Further details are provided in the Supplementary Note.

### Replication data sets

We replicated key genetic associations in the BioBank Japan (BBJ^31^) and *All of Us* (AoU^47^) cohorts. For rare pLoF CNV associations with height, we performed replication analyses in BBJ (*n*=179,420) using a SNP-array-based CNV call set we previously generated^9^. For associations with hypertension (at *RGL3*) and type 2 diabetes (at *RASA4* and *CTRB2*), we performed replication in AoU by genotyping each variant under consideration from high-coverage whole-genome sequencing data (*n*=245,394 in the AoU v7 release). Additionally, for variants with potential transcriptional effects (at *RGL3* and *SIGLEC14/SIGLEC5*), we performed follow-up in the Genotype-Tissue Expression (GTEx^33^) data set (*n*=838 in GTEx v8). Details are provided in the Supplementary Note.

### Overview of HMM method for haplotype-informed rare CNV detection

CNV-calling from exome-sequencing data typically involves searching for consistent increases or consistent decreases in a sample’s WES read coverage across a series of captured genomic regions, indicating the presence of a duplication or deletion. This requires accurately modeling WES read coverage, which can be substantially influenced by technical differences in exome capture that may vary across samples and across genomic regions (e.g., due to heterogenous effects of local GC content). While exome sequencing of UK Biobank was performed relatively uniformly across samples, exome capture was performed using a different IDT oligo lot for the first ∼50,000 samples^51^ versus the remainder of the cohort, requiring careful treatment of this batch covariate.

Our overall strategy to account for technical variation in WES read coverage (both across and within oligo lots) was to estimate sample-specific baseline models of expected read depth by identifying sets of reference samples with best-matching exome-wide coverage profiles^21^. We analyzed WES read coverage at the resolution of 100bp bins, restricting to bins with coverage in both oligo lots, similar coverage across the two oligo lots, and sufficient mappability (requiring most aligned reads to have positive mapping quality). To optimize for robust analysis of rare CNVs, we further restricted to bins in which we could accurately calibrate normalized read coverage to absolute copy number (either because a bin was rarely affected by copy-number polymorphism or because discrete copy-number states could be confidently identified).

While most WES-based CNV-callers analyze each sample independently after performing normalization, we reasoned that we could increase CNV detection sensitivity by integrating WES data across individuals likely to have co-inherited a large genomic tract (as in our recent SNP-array-based CNV analysis^9^). Similar to our previous work, we used a hidden Markov model (HMM) to call CNVs in this haplotype-informed way, integrating information regarding copy number state across an individual and up to 10 “haplotype neighbors” with expected time to most recent common ancestor (TMRCA) less than a selected value (equivalently, if the length of IBD sharing exceeded a threshold).

In more detail, for each 100bp bin, for the individual and each haplotype neighbor, we used negative binomial distributions with sample-and region-specific parameters to estimate a Bayes factor for deletion and duplication states based on counts of read alignments within the 100bp bin for each sample. For a given threshold on minimum length of IBD sharing, we computed a haplotype-informed combined Bayes factor by multiplying Bayes factors across the target individual and all haplotype neighbors with IBD sharing passing the threshold. We ran this analysis using a set of different IBD length thresholds (trading off sensitivity to more recent vs. older mutations) and compiled calls made across these IBD parameter values.

To obtain a high-quality CNV call set, we performed subsequent filtering of various classes of calls that tended to be of lower quality (based on inspection of WES and WGS read alignments at initial calls in a pilot analysis). We also removed individuals with more than 300 CNV calls. For downstream association analyses, we masked calls on any chromosome in which we had previously called a mosaic CNV^48^. Further methodological details are available in the Supplementary Note.

### Validation of HMM-based CNV call set

To benchmark precision of the HMM-based CNV call set, we analyzed independent WGS data for 100 individuals. For each of these individuals, we assessed whether or not WGS read depth was higher (respectively, lower) than expected within the putative duplications (respectively, deletions) called. We estimated validation rate as the difference between the fraction of calls with WGS read depth in the correct direction versus the opposite direction (reasoning that false positive calls should be equally likely to have WGS read depth in either direction by chance). We also determined precision and recall for the *CCNF* exon 3 duplication and *RGL3* exon 6 partial deletion by directly genotyping these CNVs using discordant-read and breakpoint-based analyses, respectively (Supplementary Note).

### Overview of haplotype-informed analysis of common copy-altering structural variants

The HMM pipeline above was designed primarily to robustly call rare CNVs from exome-sequencing data. Hidden Markov model approaches for this task directly model read coverage generated from discrete copy-number states, which can aid statistical power and breakpoint precision when the model is accurate. However, such approaches can produce suboptimal performance when model assumptions are violated. Model violations are especially prone to occur at common CNV loci (where calibration of read-depth to copy number states can be challenging) and in segmental duplications (where high sequence homology can influence read-mapping in ways that cause copy-number alterations to have unexpected effects on read depth, and loci may contain multiple complex SVs). For these reasons, genotyping common CNVs from short read-sequencing, especially within segmental duplications, is technically challenging and requires careful modeling^52^, such that general-purpose SV analysis pipelines deployed at scale have had limited ability to assess such variation^3, 24^.

Despite these challenges, short-read sequencing read-depth, including from WES, does contain useful signatures of common copy-altering SVs. We reasoned that even if precisely characterizing such SVs from WES is intractable, the signals of copy-number variation contained in WES read-depth data could still provide approximations of structural variation that, while rough, could enable discovery of SV loci associated with phenotypes—after which the SVs involved could be precisely resolved through follow-up analyses of WGS or long-read data. We therefore developed a complementary analysis pipeline to roughly estimate copy-number variation across individuals (measured on a continuous rather than discrete scale) from WES read coverage at a broad set of predefined genomic regions (including 100bp bins, exons, and previously-reported CNVs), extending methods we recently developed to analyze variable-number tandem repeats^21^.

For each genomic region under consideration, we counted WES reads aligned to the segment and normalized these read counts using sample-specific reference panels with matched exome-wide coverage profiles (as in the first step of the HMM pipeline). Unlike the HMM pipeline, we considered low-mappability regions (e.g., within segmental duplications), and we generated two read-count measurements per region, one counting all reads regardless of mapping quality and the other counting only reads with positive mapping quality. We then evaluated which of these measurements appeared to be heritable, potentially reflecting common copy number variation in the region. To do so, we computed mid-parent vs. child correlations of normalized read counts in 704 trios, restricting further analysis to 100bp bins and exons with significant correlation and all previously-reported CNVs.

For each WES read-count measurement that potentially represented common copy number variation in a region, we used long shared SNP haplotypes to statistically phase (and simultaneously denoise) the values measured across UKB WES samples and also impute into individuals without WES data. To do so, we adapted the computational methods we previously used to analyze VNTRs^21^, improving scalability by using the positional Burrows-Wheeler transform (PBWT^53^) to rapidly identify shared haplotypes. To catch instances in which exome capture bias rather than copy-number variation was responsible for heritable variation in WES coverage (e.g., short haplotypes containing several SNPs co-located within a few hundred base pairs that influence capture efficiency), we restricted to regions for which WES and WGS read-depth measurements exhibited consistent signal. Further details are available in the Supplementary Note.

### Overview of haplotype-informed analysis of paralogous sequence variants

Beyond measuring copy number at polymorphic segmental duplications, our computational approach also enabled analysis of paralogous sequence variants (PSVs) within such segments: i.e., SNPs and indels present on varying numbers of copies of a repeated segment within a single allele. To do so, we first roughly estimated PSV genotypes from counts of WGS read alignments supporting each base (i.e., “pileups”) and then adapted our haplotype-informed approach to denoise PSV genotypes and impute them into the remainder of the UKB cohort (Supplementary Fig. 4).

In more detail, for a repeat segment of interest, we first identified all regions in the GRCh38 reference sequence paralogous to the repeat segment and extracted all WGS reads aligning to these regions. We then realigned these reads to a new reference containing only one copy of the repeat segment (plus a small buffer sequence containing the beginning of a second copy of the same repeat), facilitating harmonized ascertainment and genotyping of all common PSVs. Specifically, we estimated PSV allele fractions (PSVAF; i.e., the fraction of repeat units containing a given PSV) from pileup counts, which we then converted to absolute estimates of PSV copy number (PSVCN; i.e., the number of repeat units containing a given PSV) by scaling an individual’s total repeat copy number by PSVAF. We then used long shared SNP haplotypes to phase PSVCN and impute into individuals without WGS data using the same approach we used to analyze common copy-altering SVs. Further details are provided in the Supplementary Note.

### Association testing and statistical fine-mapping

We performed SV-phenotype association analyses on three classes of CNVs derived from the HMM-based CNV call set (defined based on (i) 100bp-bin overlap, (ii) gene overlap, and (iii) gene pLoF burden) as well as the continuous-valued estimates of common copy-number variation derived from heritable WES coverage (Supplementary Fig. 1).

We conducted association tests for our primary set of 57 quantitative traits using BOLT-LMM^25, 26^ with assessment center, genotyping array, WES release (50K / 200K / 454K / 470K / none), sex, age, age squared, and 20 genetic principal components included as covariates. We fit the linear mixed model on SNP-array-genotyped autosomal variants with MAF > 10^-4^ and missingness < 0.1 and computed association test statistics for SV genotypes defined above; a similar pipeline produced association test statistics for SNP and indel variants imputed by UK Biobank (the imp_v3 release^22^). We included all participants with non-missing phenotypes in the QC-ed European-ancestry call set described above. We removed associations potentially explainable by linkage disequilibrium (LD) with imputed SNPs and indels within 3Mb^9, 54^ (see Supplementary Note).

The associations that survived this filtering represented structural variants likely to causally influence phenotypes. We annotated CNVs on this list as syndromic based on a previously-curated list of pathogenic CNVs^55^. For associations of particular interest that arose from analysis of common copy-altering SVs, we undertook follow-up in UKB WGS or HPRC long-read assemblies^5^ to more precisely resolve SVs, after which we refined SV genotypes (using optimized analyses of UKB WES or WGS data) and undertook PSV analyses as necessary. Further details on filtering of associations and follow-up analyses at loci of interest are provided in the Supplementary Note.

### Data availability

Individual-level CNV calls and continuous-valued estimates of relative copy number in UKB will be returned to UK Biobank. Summary statistics for CNV-phenotype association tests are available at https://data.broadinstitute.org/lohlab/UKB_WES_CNV_sumstats/. Access to the following data resources used in this study is obtained by application: UK Biobank (http://www.ukbiobank.ac.uk/), BioBank Japan (https://biobankjp.org/en/), *All of Us* (https://allofus.nih.gov/), GTEx (via dbGaP, https://www.ncbi.nlm.nih.gov/gap/, accession phs000424.v8.p2).

### Code availability

Custom code used to perform haplotype-informed CNV analysis of UKB WES data is provided at https://data.broadinstitute.org/lohlab/UKB_WES_CNVs_code.tar.gz for review and will be deposited in a DOI-minted repository prior to publication.

## Notes

### Competing Interest Statement

The authors have declared no competing interest.

