## Supplementary Data for "Hidden protein-altering variants influence diverse human phenotypes"

**10 Association tests with disease phenotypes** **37**

**Supplementary Figures** **38**

**Supplementary Tables** **43**

**References** **50**

### Supplementary Notes

#### 1 Exome-sequencing read counting and normalization

Computational methods for CNV-calling from whole-exome sequencing (WES) data generally search for local deviations in sequencing read-depth, as the proportion of the genome captured by WES is too sparse to provide breakpoint-based evidence (e.g., discordant read pairs and split reads) for most CNVs [1]. We likewise implemented a read-depth approach, obtaining additional statistical resolution by incorporating information across individuals sharing extended SNP-haplotypes. In this section we provide details of how we counted WES reads and normalized read counts across samples.

##### 1.1 Read-counting in 100bp bins

We expected that our haplotype-informed approach would enable modeling WES read-depth at the resolution of 100bp (rather than the exon-level resolution of most WES-based CNV callers), so we began by partitioning the genome into 100bp bins and identifying a set of 100bp bins with WES coverage and mappability suitable for analysis. To assess WES coverage across the genome for each of the two oligo lots used in UK Biobank exome sequencing [2,3], we randomly selected 100 representative samples for each oligo lot and counted the number of reads aligned to each 100bp bin of the GRCh38 reference (counting only one read from each mate pair as detailed below).

This pilot analysis identified an initial set of 831,966 autosomal 100bp bins with a mean read count of at least 5 in the second oligo lot (used to sequence all but the first $\sim$50K WES samples). For hidden Markov model (HMM)-based CNV-calling, we then filtered to a subset of 100bp bins that satisfied the following additional conditions:

- Mean read count at least 5 in both oligo lots, restricting to uniquely-mapped reads (i.e., read with nonzero mapping quality).
- Similar coverage in both oligo lots (i.e., mean uniquely-mapped read count agreeing within a factor of 2).
- Most reads (>90%) uniquely mapped (based on pilot samples from the second oligo lot).

We applied these more-stringent filters to the 100bp bins to be analyzed in HMM-based CNV-calling to minimize the potential for model violation due to technical artifacts. We reasoned that we could optimize the HMM pipeline for robustly calling rare CNVs, leaving broader analysis of common CNVs in more difficult regions (e.g., segmental duplications with low-mappability) for a separate analysis pipeline (Section 6). To further filter 100bp bins for mappability, we applied a final check to eliminate 100bp bins that represented ”mappability islands” (i.e., 100bp bins flanked by bins that failed the >90% unique-mapping criterion, or pairs of consecutive 100bp bins flanked by pairs of bins that failed the >90% unique-mapping criterion). These filters resulted in a final set of 720,151 autosomal 100bp bins for HMM-based CNV-calling.

After selecting this set of 100bp bins, we performed read-counting on each UKB WES OQFE CRAM file in turn. We counted reads that satisfied the following criteria:

- SAM flag 81, 83, 145, or 147 (i.e., read reverse strand, mate forward strand, usually corresponding to the right-hand read in a pair) [4].
- Read and mate aligned to the same chromosome.
- Read uniquely mapped (MAPQ>0).

We counted each such read as belonging to the 100bp bin containing its leftmost aligned base. This procedure ensured that each read pair (corresponding to a sequenced DNA fragment) was counted at most once, allowing read counts from different 100bp bins to be subsequently analyzed as independent measurements.

#### 1.2 Normalization using sample-specific reference sets

Counts of reads generated from copy-number-polymorphic regions are expected to scale linearly with copy number and with sequencing coverage, such that naively, information about copy number of a genomic region can be obtained simply by normalizing read counts by overall sequencing coverage (per-sample). However, variation in exome-sequencing coverage across the genome can also be strongly influenced by technical "batch effect" biases that vary across samples. To account for such effects, we applied a normalization approach in which, for each sample, we identified a set of 300 other samples that exhibited similar exome-wide coverage profiles and could thus serve as a reference set likely to be well-matched for any technical biases. We previously applied this approach in the initial UKB ∼50K WES release to normalize WES read counts for the purpose of measuring VNTR lengths [5]; here we used the same method with a few minor modifications.

Specifically, we generate a coverage profile for each WES sample at 1kb resolution by aggregating 100bp-bin read counts in 1kb windows of the genome, excluding windows that intersected known structural variant polymorphisms (SVs with MAF>0.1% in the 1000 Genomes Phase 3 call set [6], CNVs with MAF>0.1% in the 1000 Genomes high-coverage call set [7], and 118 coding VNTRs that we previously analyzed [5]). We further restricted to 1kb windows with mean WES read depth between 5–100x. We then normalized and further filtered these coverage profiles as previously described [5], with the slight modification that we cropped standardized read-depth measurements to a $z$-score range of $\pm 3$ (rather than $\pm 2$).

Finally, for each sample, we selected its 300 reference samples to be those with best-matching final coverage profiles (based on the sum of squared differences in transformed read depths across the final filtered set of 1kb windows). We performed reference-identification within WES release batches (i.e., for samples progressively added in the 50K, 200K, 454K, and 470K UKB WES data releases, we selected reference samples from those sequenced in the same release). Additionally, to reduce the potential for ancestry or relatedness to influence selection of reference samples (due to the possibility of coverage profiles still partially reflecting true copy-number variation, despite our exclusion of known CNV regions), we restricted reference samples to the 409K White British ancestry subset and excluded related samples [8].

##### 1.3 Output of read-counting and normalization

This procedure resulted in the following measurements per WES sample:

- Observed read counts (vector of 720,151 measurements; 1 per 100bp bin)
- Coverage (scalar; exome-wide coverage factor)
- Baseline read counts (vector; average across 300 reference samples of observed read counts / coverage)
- Expected read counts (vector; baseline read counts $\times$ coverage)
- Normalized read counts (vector; observed read counts / expected read counts)

We performed some subsequent modeling on normalized read counts (which are continuous-valued and centered at 1) and other modeling on observed read counts (which are discrete); in the latter case, we used the quantities derived above to model the distribution from which observed read counts were drawn. Details are provided in the following sections.

#### 2 Modeling exome-sequencing read counts

To model WES read counts within an HMM-based CNV-calling pipeline, we needed to estimate, for each sample and each 100bp bin, the distribution of read counts that would be generated from each copy-number state (CN = 1, 2, 3). We elected to model observed WES read counts using negative binomial distributions with sample- and bin-specific parameters which we will now describe.

##### 2.1 Negative binomial model

The negative binomial distribution is equivalent to a continuous mixture of Poisson distributions Poisson($\lambda$) in which the rate parameter $\lambda$ is a random variable with a gamma distribution. This mixture distribution is often used to model count data, including in WES-based CNV callers [1], as the gamma distribution naturally captures variability in the rate parameter (e.g., from unknown batch effects) that causes the count distribution to be overdispersed relative to a Poisson distribution with fixed rate parameter.

Explicitly, we assume observed read counts follow a negative binomial distribution $\mathrm{NB}(r, p)$ with parameters $r = k$ and $p = \frac{1}{1+\theta}$, where $k$ and $\theta$ are the shape and scale parameters of a gamma distribution. This is equivalent to assuming that read counts are drawn from a Poisson distribution with $\lambda \sim \mathrm{Gamma}(k, \theta)$. (Note that these formulas assume the negative binomial parameterization $X \sim \mathrm{NB}(r, p)$ in which $\Pr(X = k) = \binom{k+r-1}{k} p^r (1-p)^k$ (where $k$ is a dummy variable here, not to be confused with the shape parameter of the gamma); references differ on how the negative binomial distribution is parameterized.)

The gamma distribution $\mathrm{Gamma}(k, \theta)$ has property that scaling the distribution by a factor of $c$ corresponds to multiplying the scale parameter $\theta$ by $c$: that is, if $X \sim \mathrm{Gamma}(k, \theta)$, then $cX \sim \mathrm{Gamma}(k, c\theta)$. Additionally, the mean of the gamma distribution $\mathrm{Gamma}(k, \theta)$ is $k\theta$, such that the shape parameter $k$ can then be thought of as modulating the amount of overdispersion in the gamma-Poisson (equivalently, negative binomial) distribution.

This intuition leads naturally to a model in which for a given 100bp bin, we assume that the gamma distribution underlying the rate parameter for a given sample $i$'s Poisson draw has scale parameter $\theta_i$ satisfying:

$$\theta_i = (\text{sample } i\text{'s coverage}) \times (\text{sample } i\text{'s bin-level baseline}) \times \frac{\text{sample } i\text{'s CN}}{2} \times \theta_0, \qquad (1)$$

reflecting three scaling factors: (i) the sample's exome-wide read coverage, (ii) its "baseline" for the bin (reflecting batch effects that might cause a sample's expected coverage-adjusted read counts in a bin to be lower/higher than average, as defined in Section 1.3), and (iii) its copy number at the bin (e.g., scaling by 0.5 for DEL and 1.5 for DUP). The remaining term $\theta_0$ is a bin-specific constant that needs to be estimated together with the shape parameter of the gamma distribution ($k$, also bin-specific).

#### 2.2 Estimating bin-specific negative binomial parameters

Because the mean of a gamma distribution is given by the product of its shape and scale parameters, we can write (for a given bin):

$$\begin{aligned} &\mathbb{E}(\text{sample } i\text{'s read count}) \\ &\quad = \mathbb{E}(k\theta_i) \\ &\quad = k \times \underbrace{(\text{sample } i\text{'s coverage}) \times (\text{sample } i\text{'s baseline})}_{\text{sample } i\text{'s expected read count}} \times \frac{\mathbb{E}(\text{sample } i\text{'s CN})}{2} \times \theta_0. \end{aligned}$$

Note that on the left, the expectation is over both sampling from the gamma-Poisson and over the population distribution of copy number (CN), whereas on the right, the expectation is simply over the population distribution of CN. Canceling terms, we obtain the relationship

$$k\theta_0 = \frac{2}{\mathbb{E}(CN)}$$

providing one constraint on the parameters $k$ and $\theta_0$ to be estimated.

For a bin that does not overlap any common CNVs, we can further assume that most individuals in the population have copy number 2, such that we obtain the constraint $k\theta_0 = 1$, leaving a one-dimensional parameter optimization (say, over $\theta_0$) to obtain a maximum likelihood estimate:

$$\hat{\theta}_0 = \underset{\theta_0}{\operatorname{argmax}} \prod_i \mathcal{L}(\text{observed read count for sample } i). \tag{2}$$

To simplify this optimization, we further set the likelihood for each sample $i$ to the negative binomial likelihood corresponding to CN=2: that is, we assume that sample $i$'s read count is drawn from a single gamma-Poisson with shape parameter $k = 1/\theta_0$ and scale parameter

$$\theta_i = (\text{sample } i\text{'s coverage}) \times (\text{sample } i\text{'s baseline}) \times \theta_0,$$

obtained by setting CN=2 in equation (1).

The CN=2 assumption is of course violated when some samples have a CNV overlapping the bin. However, for a bin that only overlaps rare CNVs—such that for a small fraction of samples, read counts are likely to be too low (for DELs) or too high (for DUPs)—this model violation should only lead to a slight overestimate of overdispersion, ultimately producing slightly conservative CNV calls, which we consider acceptable. For bins that overlap common CNVs, we used a separate parameter-estimation approach detailed in Section 2.3 below.

Computationally, we performed the one-dimensional likelihood maximization in equation (2) using quadratic iteration over the range $10^{-4} \leq \theta_0 \leq 1$ starting with an initial guess $\theta_0$ based on matching first and second moments. Also, rather than performing this optimization on the full UKB WES cohort all at once, we split the computation into 16 batches (subdividing WES releases), such that for each bin, we ended up with 16 estimated values of $\theta_0$ and $k$, one per sample batch. This procedure was computationally convenient and also handled the difference in oligo lots used for the first 50K WES samples versus subsequence WES releases.

#### 2.3 Estimating bin-specific negative binomial parameters in common CNV regions

As noted in the previous section, modeling sequencing read counts in common CNV regions is more challenging because observations from different samples derive from different distributions (due to copy number variation across samples). In particular, in regions with lower signal-to-noise, the task of simultaneously estimating model parameters and population frequencies of CNVs is challenging and requires careful analysis [9, 10].

To enable inclusion of such regions in our HMM-based CNV-calling pipeline, we used a separate approach to estimate negative binomial parameters for 100bp bins overlapping common CNVs. This approach involved (1) identifying bins with common copy-number variation, (2) identifying copy-number modes in normalized read counts measurements, and (3) estimating negative binomial parameters derived from the inferred modes. We found it convenient to work with continuous-valued normalized read counts (i.e., observed divided by expected read counts, based on a sample's coverage and baseline; Section 1.3) in the first two steps before converting back to the negative binomial formulation in the third step.

##### 2.3.1 Identifying bins with common copy-number variation

To identify 100bp bins likely to be commonly copy-number-polymorphic, we searched for heritable variation in normalized read counts from 704 exome-sequenced trios in UK Biobank. That is, for each bin, we computed the correlation between normalized read counts in children versus parental averages, reasoning that common copy-number variation should cause these values to be positively correlated within trios. We identified bins that required further modeling as those for which the correlation was statistically significant ($P < 0.01$) and had magnitude at least what would be expected for a deletion with 5% allele frequency (taking into account each bin's coefficient of variation of normalized read depth), reasoning that bins overlapping only rarer CNVs are reasonably well-modeled using the approach of Section 2.2.

##### 2.3.2 Identifying copy-number modes in normalized read counts

For each bin exhibiting heritable read-depth according to the above procedure, we performed further analysis of the distribution of normalized read counts across samples to estimate the copy-number distribution in the population (described in this section) and then to estimate negative binomial parameters (described in Section 2.3.3 below). We performed these analyses once per oligo lot using the highest-coverage 1,000 samples from each oligo lot in the 200K WES release, reasoning that common copy-number modes would be easier to identify in a subset of less-noisy samples.

**Constrained Gaussian mixture model.** Following previous work (e.g., Genome STRiP [9] and CLAMMS [10]), we attempted to fit the distribution of normalized read counts across samples (for a given bin under consideration) using three possible constrained Gaussian mixture models:

- DEL-only. Letting $p$ denote the allele frequency of the DEL (assumed to be in Hardy-Weinberg equilibrium), this model comprises a mixture of three Gaussians corresponding to

CN=$\{0, 1, 2\}$ with mixture weights $\{p^2, 2p(1-p), (1-p)^2\}$, means $\{0, 0.5\mu, \mu\}$, and variances $\{0.5\sigma^2, 0.5\sigma^2, \sigma^2\}$. (The variance of the CN=0 mode is set to be nonzero to improve robustness, even though in theory no reads should be observed.) Assuming that the mean of the mixture distribution is 1 (due to the normalization of read counts), we have $\mu = \frac{1}{1-p}$, leaving two parameters to optimize: $\sigma^2$ and $p$.

- DUP-only. Letting $p$ denote the allele frequency of the DUP, this model comprises a mixture of three Gaussians corresponding to CN=$\{2, 3, 4\}$ with mixture weights $\{(1-p)^2, 2p(1-p), p^2\}$, means $\{\mu, 1.5\mu, 2\mu\}$, and variances $\{\sigma^2, 1.5\sigma^2, 2\sigma^2\}$. Assuming again that the mean of the mixture distribution is 1, we have $\mu = \frac{1}{1+p}$, again leaving $\sigma^2$ and $p$ to optimize.

- DEL+DUP. Letting $\alpha$ denote the allele frequency of the DEL and $\beta$ denote the allele frequency of the DUP, this model comprises a mixture of five Gaussians corresponding to CN=$\{0, 1, 2, 3, 4\}$ with mixture weights $\{\alpha^2, 2\alpha(1-\alpha-\beta), 2\alpha\beta+(1-\alpha-\beta)^2, 2\beta(1-\alpha-\beta), \beta^2\}$, means $\{0, 0.5\mu, \mu, 1.5\mu, 2\mu\}$, and variances $\{0.5\sigma^2, 0.5\sigma^2, \sigma^2, 1.5\sigma^2, 2\sigma^2\}$. Assuming again that the mean of the mixture distribution is 1, we have $\mu = \frac{1}{1-\alpha+\beta}$, leaving three parameters to optimize: $\sigma^2$, $\alpha$, and $\beta$.

We fit these constrained Gaussian mixture distributions imposing the following constraints on the parameters being optimized ($(\sigma^2, p)$ or $(\sigma^2, \alpha, \beta)$):

- $0 < \sigma^2 < \hat{\sigma}^2$ (fitted variance must be less than observed variance)
- $0 < p < 1$
- $0 < \alpha, \beta < 1$
- $\alpha + \beta < 0.75$ (allele frequency of REF allele with haploid CN=1 must be >25%)

**Evaluating goodness of fit.** We required reasonable similarity between fitted and observed distributions of normalized read counts (which also served as a check for consistency of observed data with Hardy–Weinberg equilibrium, as the fitted models assume Hardy–Weinberg equilibrium). For each of the constrained DEL, DUP, and DEL+DUP model fits, we measured similarity between the observed distribution and random samples from the fitted distribution using optimal transport.

In more detail, for each constrained model fit, we generated a random sample of the same size as the observed sample. Additionally, for the DEL and DEL+DUP models, we generated a random sample in which samples from the CN=0 state were forced to be exactly zero. For each of these five random samples, we sorted the sampled values, and we then computed the mean difference between each sampled value and the corresponding value in the sorted list of observed data points. We considered the DUP model to be deviant if this mean difference exceeded 0.05, and for the DEL and DEL+DUP models, we considered their fits to be deviant if both versions of the random sample (with and without setting CN=0 samples to zero) failed the 0.05 similarity threshold.

**Model selection.** We used a likelihood-ratio test to compare each model that passed the similarity-based goodness-of-fit check (DEL, DUP, and/or DEL+DUP) to a null model assuming no CNV (in which normalized read counts are drawn from a single Gaussian with mean of 1 and the observed standard deviation). We computed $P$-values from the likelihood-ratio test statistic for each model using the appropriate $\chi^2$ distribution (1-df for the DEL and DUP models and 2-df for the DEL+DUP model) and identified the model with the lowest $P$-value.

To restrict HMM-based analysis of common CNV regions to those with high-confidence model fits, we retained a common-CNV 100bp bin for further analysis only if its best model achieved $P < 10^{-25}$. More precisely, given that we ran all of the above modeling independently on high-coverage samples from each of the two oligo lots, we retained a bin for a given oligo lot if either (a) its best model in that oligo lot achieved $P < 10^{-25}$, or (b) its best model (DEL, DUP, or DEL+DUP) agreed with the best model achieving $P < 10^{-25}$ in the other oligo lot (i.e., we allowed a high-confidence fit in one oligo lot to "rescue" a lower-confidence fit in the other).

Models selected in this way were taken forward for incorporation into the negative binomial framework for directly modeling integer read counts (described next), while common-CNV bins without a confident model fit for a given oligo lot were excluded from further analysis (for samples sequenced with that oligo lot).

##### 2.3.3 Estimating negative binomial parameters using method of moments

For 100bp bins for which normalized read counts were well-fit by a constrained Gaussian mixture model according to the above criteria, we estimated parameters for our negative binomial model of discrete read counts using the method of moments. That is, we matched first and second moments of the CN=2 component of the fitted Gaussian mixture (i.e., $\mu$ and $\sigma^2$ in the notation of Section 2.3.2) to the corresponding moments that would be produced under the negative binomial model assuming CN=2. Explicitly:

$$\begin{aligned}
\mu &= \mathbb{E}(\text{normalized read counts} \mid \text{CN} = 2) \\
&= \frac{1}{n} \sum_i \mathbb{E}(\text{sample } i\text{'s normalized read counts} \mid \text{CN} = 2) \\
&= \frac{1}{n} \sum_i \mathbb{E}\left( \frac{\text{sample } i\text{'s read counts}}{\text{coverage}_i \times \text{baseline}_i} \mid \text{CN} = 2 \right) \\
&= \frac{1}{n} \sum_i \frac{k\theta_{i|\text{CN}=2}}{\text{coverage}_i \times \text{baseline}_i} \\
&= \frac{1}{n} \sum_i \frac{k \times \text{coverage}_i \times \text{baseline}_i \times \theta_0}{\text{coverage}_i \times \text{baseline}_i} = k\theta_0
\end{aligned}$$

and similarly,

$$\begin{aligned}
\sigma^2 &= \text{Var}(\text{normalized read counts} \mid \text{CN} = 2) \\
&= \frac{1}{n}\sum_i \text{Var}(\text{sample } i\text{'s normalized read counts} \mid \text{CN} = 2) \\
&= \frac{1}{n}\sum_i \text{Var}\left(\frac{\text{sample } i\text{'s read counts}}{\text{coverage}_i \times \text{baseline}_i} \mid \text{CN} = 2\right) \\
&= \frac{1}{n}\sum_i \frac{k\theta_{i|\text{CN}=2}(1+\theta_{i|\text{CN}=2})}{(\text{coverage}_i \times \text{baseline}_i)^2} \\
&= \frac{1}{n}\sum_i \frac{k\theta_0 \times \text{coverage}_i \times \text{baseline}_i(1+\theta_0 \times \text{coverage}_i \times \text{baseline}_i)}{(\text{coverage}_i \times \text{baseline}_i)^2} \\
&= \frac{1}{n}\sum_i \frac{k\theta_0(1+\theta_0 \times \text{coverage}_i \times \text{baseline}_i)}{\text{coverage}_i \times \text{baseline}_i} \\
&= \left(\frac{1}{n}\sum_i \frac{k\theta_0}{\text{coverage}_i \times \text{baseline}_i}\right) + k \times \theta_0^2 \\
&= k\theta_0\left(\frac{1}{n}\sum_i \frac{1}{\text{coverage}_i \times \text{baseline}_i}\right) + k\theta_0^2.
\end{aligned}$$

That is, we have:

$$\begin{aligned}
\mu &= k\theta_0 \\
\sigma^2 &= k\theta_0\left(\frac{1}{n}\sum_i \frac{1}{\text{coverage}_i \times \text{baseline}_i}\right) + k\theta_0^2.
\end{aligned}$$

The term $\frac{1}{n}\sum_i \frac{1}{\text{coverage}_i \times \text{baseline}_i}$ is an easily computed constant, such that the two equations above can be solved for $k$ and $\theta_0$ given $\mu$ and $\sigma^2$ from the CN=2 component of the selected Gaussian mixture model:

$$k = \frac{\mu}{\theta_0}, \quad \theta_0 = \frac{\sigma^2}{\mu} - \left(\frac{1}{n}\sum_i \frac{1}{\text{coverage}_i \times \text{baseline}_i}\right).$$

Finally, because moment-based estimation can produce out-of-range estimates (e.g., negative values of $\theta_0$), we lower-bounded $\theta_0$ at $10^{-4}$ (corresponding to minimal overdispersion).

#### 2.4 Estimating sample-specific negative binomial parameter adjustments

The bin-specific parameters estimated using the above procedures accounted for bin-level variation in the extent to which 100bp bins exhibited overdispersion. We implemented one further adjustment to the negative binomial parameters to account for sample-level variation in overdispersion—i.e., the extent to which some samples had noisier WES read counts than others (based on the behavior of read counts from a given sample across all modeled 100bp bins). Specifically, for each sample, we estimated sample-specific adjustment factors that we multiplied into the shape parameter $k$ and divided from the scale parameter $\theta_0$ of each bin, selecting these adjustment factors to improve the overall likelihood of the sample's observed read counts. We noticed that the appropriate adjustment seemed to differ somewhat for bins with lower versus higher average coverage, so we partitioned the set of 100bp bins into coverage deciles, after which we estimated a sample-specific adjustment factor for each decile by maximizing likelihood of observed read counts for bins in that decile.

### 3 Haplotype-informed CNV-calling from WES read counts

Here we describe the hidden Markov model (HMM) algorithm that we used to call CNVs by integrating probabilistic information about copy-number state derived from WES read counts of individuals and their "haplotype neighbors" (i.e., other individuals in the cohort sharing long SNP-haplotypes).

#### 3.1 Estimating per-bin Bayes factors for copy-number states

For each sample, for each 100bp bin, we used the negative binomial model parameters estimated above (Section 2) to quantify the extent to which a given read count measurement supported the presence of a copy-gain, copy-loss, or no CNV spanning the bin. Explicitly, we took the estimated bin-specific shape ($k$) and scale ($\theta_0$) parameters (from Section 2.2 for non-common-CNV bins and Section 2.3 for common-CNV bins), applied the estimated sample-specific adjustments to these parameters (from Section 2.4), and further scaled the scale parameter by the sample's coverage, bin-specific baseline, and copy-number state (CN=1,2,3) divided by 2 (equation (1)). The negative binomial distribution $\mathrm{NB}(r = k, p = \frac{1}{1+\theta})$ then gave the likelihood of the observed read count assuming each copy-number state, from which we estimated Bayes factors for copy-gain vs. no CNV and for copy-loss vs. no CNV. (We did not explicitly include CN=0 or CN=4 states in the HMM, reasoning that read counts produced by CN=0 (resp. CN=4) would heavily favor copy-loss (resp. copy-gain) within a model including CN=1,2,3.) We cropped Bayes factors to the range $[10^{-4}, 10^4]$ to limit the influence of potential outlier values.

**Filters on read count measurements.** To improve robustness of CNV-calls, we additionally implemented several filters on read count measurements considered in analysis (beyond the bin-level filters described in the previous sections):

- Additional bin-level filters:
  - For each non-common-CNV bin, we required that the coefficient of variation across samples of estimated baseline read counts for that bin (as defined in Section 1.3) was not excessively high (i.e., $<0.2$ in the majority of the 16 sample batches on which we performed read count-modeling). This filter eliminated a small number of 100bp bins for which read counts appeared to be strongly affected by technical variation.
  - For each common-CNV bin with a high-confidence constrained Gaussian mixture model fit for both oligo lots, we required the means of the CN=2 component estimated in the two oligo lots to agree within 5%.
- Additional read count measurement-level filters:
  - For each sample and each bin, we required that the estimated baseline read count (as defined in Section 1.3) did not differ excessively from the mean baseline for that bin (i.e., at least 4/5 and no more than 4/3 of the mean baseline). This filter protected against the possibility of a sample's reference set being mismatched, such that normalizing using the reference set (which intuitively corresponds to dividing by the baseline) could never shift normalized read counts by more than a factor of 0.75–1.25, i.e., halfway to a DEL or DUP.
  - For each sample and each bin, we further required that the 300bp region containing the bin and its 100bp flanks not contain too many nearby SNPs (i.e., 4 or more ALT genotype calls at REF-major positions or 4 or more ALT genotype calls at REF-minor positions). This filter protected against potential exome capture bias produced by polymorphisms that might interfere with oligo binding.

#### 3.2 Hidden Markov model (HMM) for CNV-calling

We used a three-state HMM with copy-gain, copy-loss, and no-CNV states to analyze each haplotype of each individual. For a given haplotype of a given individual, for each 100bp bin, we used the negative binomial probabilities detailed above (Section 3.1) to estimate log Bayes factors for copy-gain vs. no CNV and for copy-loss vs. no CNV from the WES read count of the individual and likewise for each "haplotype neighbor" (i.e., the top 10 other individuals in the cohort with longest IBD matches spanning the bin under consideration, which we identified as previously described [11]). As we performed inference based on the Viterbi path through the HMM, we performed all computations in log space, working only with relative emission probabilities (i.e., log Bayes factors).

To aggregate information from an individual and haplotype neighbors into a single log Bayes factor per bin for copy-gain (respectively, copy-loss) vs. no CNV, we computed a sum of the individual's log Bayes factor and the log Bayes factors of all haplotype neighbors expected to share a common ancestor within the past $T$ generations. Explicitly, as derived previously [11, 12], the probability that an IBD segment of length $l$ Morgans had a TMRCA within $T$ generations is:

$$P(t < T|l) = e^{-2lT}(1 + 2lT + \frac{1}{2}(2l)^2T^2).$$

If, for a given haplotype neighbor, $P(t < T|l) \geq 0.5$, we incorporated WES read count information from this neighbor; otherwise, we ignored this neighbor. Intuitively, this setup optimizes for detecting CNVs that arose roughly $T$ generations ago (by incorporating information from haplotype neighbors who share more recent IBD—and thus have read coverage informative of the co-inherited CNV—while discarding information from haplotype neighbors with TMRCA predating the CNV mutation). To power detection of CNVs of different ages, we ran HMM inference using six different values of $T \in \{0, 5, 10, 25, 50, 100\}$ generations, where $T = 0$ corresponds to ignoring haplotype neighbors entirely (i.e., performing single-sample analysis). For each $T > 0$, we performed two HMM runs, incorporating information from neighbors of each of the individual's two haplotypes in turn. This haplotype-informed strategy also naturally allowed us to call CNVs in the $\sim$3% of UK Biobank participants who were not exome-sequenced (by computing log Bayes factors using only exome-sequenced neighbors).

We specified a symmetric transition probability matrix in which probabilities for copy-number state changes between consecutive bins depend on the distances between the bins (such that state transitions between nearby bins are less likely than between distant bins):

| from/to | CN=1 | CN=2 | CN=3 |
|---|---|---|---|
| CN=1 | $1-p-p^2$ | $p$ | $p^2$ |
| CN=2 | $p$ | $1-2p$ | $p$ |
| CN=3 | $p^2$ | $p$ | $1-p-p^2$ |

We defined the single-copy-number transition probability $p$ as $p = a(1 - e^{-rd})$, where $a$ represents the asymptote (i.e., the probability of a transition between very distant bins, set to be $10^{-3}$), $r$ represents a relaxation rate (defining what distance scale corresponds to "very distant," set to be $10^{-3}$), and $d$ represents the base-pair distance between consecutive bins.

### 4 Merging and filtering CNV calls

We synthesized CNV calls from the HMM (generated using six different values of the TMRCA parameter $T \in \{0, 5, 10, 25, 50, 100\}$) in a manner similar to our previous haplotype-informed CNV analysis of SNP-array probe intensity data [11].

#### 4.1 Post-processing CNV calls from each HMM run

For each individual and each run of the HMM (parameterized by $T$ and by the haplotype of the individual used to identify neighbors), we extracted potential deletions (respectively, duplications) as consecutive sequences of copy-loss (respectively, copy-gain) states in the Viterbi path through the HMM. For each such sequence of states, we computed the $\log_{10}$ Bayes factor (BF) supporting the putative CNV event (as the sum of $\log_{10}$BFs across the sequence of bins within the segment, including information from the target individual as well as haplotype neighbors as in the HMM).

We post-processed the segments by bridging short gaps between consecutive segments of the same copy-number state (because the Viterbi path through long CNVs was sometimes interrupted by short sequences of no-CNV states). Specifically, we bridged gaps between nearby CNV segments if they included at most 3 bins (i.e., next start bin index – current end bin index $\leq 4$) and spanned <10 kb (i.e., next start bp – current end bp $< 10,000$).

#### 4.2 Merging and filtering CNV calls across HMM runs

To synthesize post-processed CNV calls across HMM runs from different values of the TMRCA parameter $T$ (which had differing sensitivity to CNVs of different mutational ages and also exhibited stochastic variation in endpoints), we next performed a deduplication step to identify a nonredundant set of CNVs discovered in each individual. We performed this deduplication procedure on the aggregate set of CNV calls made across values of $T$ and across which of the individual's haplotypes had been used to identify neighbors. (Homozygous CNVs present on both haplotypes were collapsed into a single call during this step.)

Specifically, we considered two CNV calls of the same type (DUP or DEL) to be duplicates if their endpoints matched within 4 bins (i.e., $\Delta$start $\leq$ 4 and $\Delta$end $\leq$ 4). For each such duplicate pair, we retained the call with higher $\log_{10}$ BF.

We then applied a set of filters to these potential CNVs. First, for each putative CNV identified for an individual $i$, we computed the contribution to the CNV's $\log_{10}$BF from only individual $i$'s WES read count data (i.e., not considering haplotype neighbors), which we denote $\log_{10}\text{BF}_{i\text{-only}}$. For deletions, we required either (i) $\log_{10}\text{BF}_{i\text{-only}} > 0$ (i.e., the individual's WES read counts support the call) or (ii) $\log_{10}\text{BF}_{i\text{-only}} = 0$ and $\geq$3.5 haplotype neighbors were used on average across bins within the call (i.e., the individual itself provides no information, typically because WES was unavailable, but several haplotype neighbors support the call). For duplications, we required that the call span at least 300bp and have $\log_{10}\text{BF}_{i\text{-only}} \geq 0$ (i.e., the individual's WES read counts do not disagree with the call). These requirements served to prevent haplotype neighbors from overruling direct evidence against a CNV from a sample's WES data.

We then applied a set of cohort-wide filters. We removed three specific calls which were made in almost every sample and likely represented technical artifacts (specifically, chr2:232,408,200–

232,409,800, chr18:11,609,900-11,610,200, and chr19:49,054,900-49,055,400; we removed any call with endpoints matching within $\pm 1$ bin). We further removed deletions $\leq$200 bp with $\leq$10,000 UK Biobank carriers and $\log_{10}\mathrm{BF}_{i\text{-only}} < 1$ (i.e., we required rare short DELs to have at least moderate support from a sample's own WES data).

Because this call set could still contain overlapping CNV calls, we next merged overlapping CNV calls of the same type (DUP or DEL). Lastly, we applied a final set of density filters on the CNV calls, removing any calls longer than 10Mb with length / number of bins $> 10^5$ (i.e., we required that extremely long calls be supported by data from at least 1 bin per 100kb).

##### 4.3 Excluding individuals with aberrantly many CNV calls

To further improve robustness of the call set, we removed individuals with $>$300 CNV calls potentially indicating technical artifacts (running this filter after excluding calls on chromosomes containing mosaic chromosomal alterations with cell fraction $>$20% [13], as large mosaic CNVs can result in many short CNVs being called across the span of the mosaic CNV).

#### 5 Properties of UK Biobank WES-derived CNV call set

##### 5.1 Annotation of gene-overlapping CNVs

**Gene annotations.** We downloaded canonical transcripts from UCSC (`http://hgdownload.soe.ucsc.edu/goldenPath/hg38/database/knownCanonical.txt.gz`). We extracted locations of CDS from GENCODE v41 transcripts (`https://ftp.ebi.ac.uk/pub/databases/gencode/Gencode_human/release_41/gencode.v41.annotation.gtf.gz`). We restricted to CDS from protein-coding canonical transcripts (available for 19,539 genes). For each canonical transcript, we extracted the start and end of each CDS as well as the start and end of the transcript.

**Genic consequences of CNVs.** We annotated CNVs with the following genic consequences:

- DUP: whole-gene duplication (starting before than the start of the transcript and ending after the end of the transcript)
- pLoF (DEL-only): deletion intersecting a CDS
- pLoF (DEL and IED): deletion intersecting a CDS or intragenic exonic duplication (IED; i.e., duplication intersecting a CDS, starting at least 1kb downstream of the start of the transcript and ending at least 1kb upstream of the end of the transcript)
- pLoF (high-confidence): $\geq$300bp deletion intersecting a CDS.

#### 5.2 Gene constraint metrics

We computed the number of UK Biobank participants of European ancestry carrying whole-gene duplications and CNVs predicted to cause loss of function for each protein-coding gene (high-confidence pLoF; see Section 5.1) and compared these DUP and pLoF frequencies to measures of gene constraint. We annotated each gene with its LOEUF sextile bin (`oe_lof_upper_bin_6` from the pLoF Metrics by Gene TSV file downloaded from `https://gnomad.broadinstitute.org/downloads` [14]), restricting to genes with a non-missing LOEUF sextile bin and genes with only one annotated canonical transcript. In Fig. 1e, we reversed the order of LOEUF sextile bins such that higher-numbered bins correspond to more-constrained genes. We also extracted pLI (probability of loss-of-function intolerance) scores from the same file.

#### 5.3 Rare gene-altering CNVs

We used the following procedure to compute numbers of rare gene-altering CNVs per individual (for the distributions plotted in Fig. 1f):

1. Restrict to genes rarely altered by either (a) pLoF (high-confidence DELs) or (b) whole-gene DUPs. We defined rare to be $<1\%$ allele frequency (i.e., $<2\%$ of individuals are carriers).
2. For each individual, we then restricted to CNVs that either (i) were annotated as high-confidence pLoF DEL for a gene rarely altered by such an event (i.e., in gene set (a) above); or (ii) were annotated as a whole-gene DUP for a gene rarely altered by such an event (i.e., in gene set (b) above).
3. For each individual, we then counted the number of CNVs of type (i) and of type (ii) as well as the number of genes affected by each type of CNV (i.e., a duplication that overlaps 2 rarely-duplicated genes would count as 1 rare whole-gene DUP called but 2 genes affected by rare whole-gene DUPs).

#### 5.4 Validation rate based on WGS read-depth analysis

To assess the precision (i.e., validation rate) of CNVs called by our HMM pipeline, we computed the proportion of calls that exhibited enrichment or depletion of WGS read-depth consistent with the CNV call. We performed these analyses using whole-genome sequencing data for 100 individuals in our primary analysis set.

To calibrate WGS read-depth measurements, which we computed using mosdepth [15], we adjusted for genome-wide sequencing depth (across autosomes). We computed mean normalized read depth across each CNV being validated. We then compared this normalized read depth to the distribution of normalized read depth across the same region among individuals with no CNV call in the region.

The above computations allowed us to classify CNV calls as either (1) supported by read-depth signal in the correct direction (e.g., depletion of read-depth for a deletion) or (2) having read-depth signal in the incorrect direction. We estimated validation rate as (1) minus (2).

### 6 Continuous-valued copy-number estimation at common structural variant loci

To enable exploration of common coding copy-number variation—including variation within segmental duplications—we developed a separate approach (complementary to the HMM pipeline described in Sections 2 and 3) that (i) identifies genomic regions that harbor common copy-altering polymorphisms and then (ii) measures copy number in these predefined regions by leveraging haplotype-sharing to denoise individual read-depth-derived estimates. The first step made use of close relatives (specifically, parent-child trios) in the UK Biobank cohort to ascertain regions that contain common copy-number variation, while the second step leveraged haplotype-sharing among the full cohort to refine copy-number measurements derived from WES read depth for all samples.

In contrast to the HMM pipeline, this common-CNV analysis approach produced continuous-valued measurements of relative copy number (rather than discrete genotype calls), broadly similar to the probabilistic dosage values typically produced by SNP imputation methods. This approach, which generalizes techniques we recently developed to study variable number tandem repeat (VNTR) polymorphisms [5], allowed more flexibility to analyze regions containing more-complex copy number variation that might be difficult to precisely model and genotype.

#### 6.1 Identifying regions harboring common copy-altering polymorphisms

To survey common copy-number variation as comprehensively as possible from exome-sequencing data, we defined a broad set of regions to examine for evidence of copy-number polymorphism, including 100bp bins, exons, and previously-reported CNVs. The rationale behind this approach was that measuring WES read coverage at finer resolutions might be necessary to capture short CNVs, whereas measuring WES read coverage across wider regions would produce the most accurate measurements of large CNVs. We therefore sought to initially measure variation at a broad set of regions (and make use of previous catalogs of common CNVs), reasoning that we could later fine-tune analyses at specific loci of interest identified from association analysis. Specifically, we considered the following types of genomic regions:

- 100bp bins with WES coverage (see Section 1.1).
- Exons. We extracted CDS regions from GENCODE v38 [16] and rounded the starting base down to the nearest 100bp and the end up to nearest 100bp.

- Previously-reported CNVs: We extracted deletions, duplications, and mCNVs that passed QC from gnomAD-SV [17] (v2.1, lifted to hg38); as well as deletions, duplications, and CNVs from the 1000 Genomes Project call sets (both the 30x-coverage ensemble SV call set [7] (freeze V1, `http://ftp.1000genomes.ebi.ac.uk/vol1/ftp/data_collections/1000G_2504_high_coverage/working/20210124.SV_Illumina_Integration/`), and the Phase 3 SV calls [6], downloaded from `http://ftp.1000genomes.ebi.ac.uk/vol1/ftp/phase3/integrated_sv_map/`). We restricted to SVs with at least one European alternate allele carrier and a minor CNF $>0.01$, and we again rounded the starting base down to the nearest 100bp and the end up to nearest 100bp.

To enable insight into regions with low-mappability, for each region, we compute two separate read-count measurements, one counting read alignments with any mapping quality (MAPQ$\geq$0; "Q0") and the other restricting to alignments with positive mapping quality (MAPQ$\geq$1; "Q1").

For each genomic region and for each mapping quality category (Q0 and Q1), we checked for evidence of common copy-number variation based on correlated coverage-normalized WES read counts among 704 parent-child trios. We took forward measurements from 100bp segments and exons (Q0 or Q1) exhibiting significant midparent-child correlation ($P < 0.01$), and we also took forward all known-CNV regions (both Q0 and Q1 measurements).

#### 6.2 Phasing and denoising normalized read counts in common-CNV regions

To efficiently phase and denoise normalized WES read counts (normalized as in Section 1.3) measuring copy number across many thousands of copy-number-polymorphic regions exome-wide, we used a simplified version of the iterative phasing algorithm we previously used to analyze VNTRs. In our previous algorithm (described in Section 3 of the Supplementary Text of ref. [5]), we updated haploid allele lengths of each individual in turn according to a probabilistic haplotype-copying model using all other haplotypes as a reference panel, prioritizing copying from haplotypes closely matching the individual's SNP-haplotypes.

Explicitly, assuming an individual's two haplotypes (carrying alleles with copy number $x_1$ and $x_2$) had been copied from reference haplotypes $i$ and $j$ with estimated allelic copy numbers $\widehat{x_i}$ and $\widehat{x_j}$, the iterative update took the form:

$$\widehat{x_1} \leftarrow \widehat{x_i} + p \cdot (\widehat{x_{1+2}} - (\widehat{x_i} + \widehat{x_j})), \qquad \widehat{x_2} \leftarrow \widehat{x_j} + p \cdot (\widehat{x_{1+2}} - (\widehat{x_i} + \widehat{x_j})) \tag{3}$$

where $\widehat{x_{1+2}}$ denotes the individual's WES-derived (diploid) copy-number estimate and $p$ denotes a constant factor between 0 and 0.5 (given by $\frac{\sigma^2_{mut}}{2\sigma^2_{mut}+\sigma^2_{err}}$ in the notation of ref. [5]). In our previous work, we considered possible updates from many pairs of reference haplotypes $i$ and $j$, and we set the updated estimates $\widehat{x_1}$ and $\widehat{x_2}$ to equal weighted averages of equation (3) across pairs

$(i, j)$, weighting contributions from different reference haplotype pairs according to their posterior probabilities of being the "source" haplotypes.

Here, we simplified the above approach by considering only 10 haplotype neighbors for each of the individual's two haplotypes and computing simple averages $\overline{\widehat{x_i}}$ and $\overline{\widehat{x_j}}$ of the copy numbers estimated across each of these sets of reference haplotypes. We then simplified the update equation to a single computation (equivalent to weighting all selected reference haplotypes equally):

$$\widehat{x_1} \leftarrow \overline{\widehat{x_i}} + p \cdot (\widehat{x_{1+2}} - (\overline{\widehat{x_i}} + \overline{\widehat{x_j}})), \qquad \widehat{x_2} \leftarrow \overline{\widehat{x_j}} + p \cdot (\widehat{x_{1+2}} - (\overline{\widehat{x_i}} + \overline{\widehat{x_j}})) \tag{4}$$

We identified haplotype neighbors using a straightforward positional Burrows-Wheeler transform (PBWT [18]) approach based on identifying longest-matching haplotype prefixes/suffixes according to adjacency within lexicographically sorted haplotype lists generated by the PBWT (run in both forward/reverse directions).

#### 6.3 Optimizing phasing parameter

The update equation (4) in our simplified phasing algorithm has a single parameter $p$ between 0 and 0.5 that determines the relative weighting of information derived from haplotype-sharing versus direct read-depth-based estimates. That is, the limit $p \to 0$ corresponds to only using information from copied haplotypes (and ignoring an individual's WES information), while the limit $p \to 0.5$ corresponds to only using an individual's WES read-depth data (and not using haplotype-sharing information at all). The optimal parameter $p$ is thus region-specific, as larger values of $p$ are suitable for regions for which WES read-depth measurements already accurately measure copy number, whereas smaller values of $p$ are suitable for regions for which WES read-depth data is very noisy, such that SNP-haplotypes must be relied upon heavily to denoise the data.

For each region (and mapping quality criterion, Q0 or Q1), we optimized the phasing parameter $p$ at the beginning of each iteration of the phasing algorithm by selecting $p$ to maximize a trio-based concordance metric. Specifically, we held out parents of UK Biobank trios from phasing, which allowed us to measure correlation between denoised estimates in children vs. averages of held-out (unphased) estimates from each pair of parents. For each value of $p$ between 0 and 0.5 in increments of 0.005, we applied the phasing algorithm with parameter $p$ to the trio children and computed this correlation. We then selected the value of $p$ that maximized trio correlation and used it to perform a full round of phasing on the full cohort (except the held-out parents). Finally, after 10 full iterations of phasing, we ran phasing on the held-out parents using the value of $p$ selected in the final iteration.

##### 6.4 Filtering regions to take forward to association analysis

We applied two additional filters to select a subset of region measurements (i.e., region and Q0/Q1 mapping quality criterion used to count reads) to take forward for further analyses. First, to reduce redundancy in the measurements (as measurements in nearby 100bp bins or exons sometimes reflected the same CNV), we performed LD-pruning using an $r^2 < 0.9$ threshold (on denoised estimates from a pilot phasing analysis of 10% of the cohort). When choosing between highly correlated regions, we retained the region with higher trio correlation of normalized WES read counts (indicating less noise in direct read-depth based estimates).

Second, to catch instances in which exome capture bias might have created heritable read-depth deviations that did not reflect copy-number variation (e.g., short haplotypes containing several SNPs within a few hundred bp that interfered with oligo binding), we filtered to regions for which WES and WGS read-depth appeared to produce consistent signal. Specifically, we computed the correlation between coverage-normalized WES read depth and WGS read depth within 500 individuals and restricted to regions with significant ($P < 0.05$) positive correlation.

#### 7 Quantitative trait association tests and statistical fine-mapping

##### 7.1 CNV genotypes and copy-number measurements tested

To search for copy-altering variants that influence phenotypes, we defined several types of genetic variables to include in association tests. From the common-CNV pipeline (Section 6), we directly tested continuous-valued copy-number measurements from normalized WES read counts (denoised by phasing). From the HMM pipeline (Sections 2 and 3), we created three categories of tests based on discrete HMM-based CNV calls:

- DELs or DUPs overlapping a given 100bp bin.
- CNVs affecting a given gene (whole-gene DUP and pLoF categories defined in Section 5.1).
- Overall pLoF burden for a given gene. Following the UK Biobank RAP documentation (`https://dnanexus.gitbook.io/uk-biobank-rap/science-corner/using-regenie-to-generate-variant-masks`), we used REGENIE [19] to build pLoF burden masks using three allele frequency thresholds (<1%, <0.1%, or singleton) for (i) pLoF SNPs/indels called from WES or (ii) pLoF SNPs/indels as well as high-confidence pLoF CNVs from our HMM pipeline (DEL and ≥300 bp).

(For the pLoF burden category, we note that the way in which allele frequency thresholds were imposed for SNPs and indels differs slightly from how they were imposed for CNVs: allele frequencies of pLoF SNPs and indels were computed for each variant separately before determining which variants met an AF cutoff, whereas a single allele frequency of pLoF CNVs was computed for each gene to decide whether to include all pLoF CNVs in the burden mask.)

Finally, for individuals with a mosaic chromosomal alteration call with cell fraction greater than 20% [13], we set all bin- and gene-level genotypes on the affected chromosome(s) to missing.

#### 7.2 Association testing

We ran BOLT-LMM [20,21] to compute linear mixed model association statistics between all types of CNV genotypes and measurements described above and 57 quantitative traits (Supplementary Data 1). To guard against potential confounding from population stratification, we restricted to individuals of self-reported White ethnicity, and we further excluded individuals with trisomy 21 or blood cancer (as described in our previous work [11]), aberrantly many CNV calls (Section 4.3), and those who had withdrawn at the time of analysis. We included as covariates assessment center, genotyping array, WES release (i.e., indicator variables corresponding to sample inclusion in the 50K, 200K, 454K, or 470K WES releases), sex, age, age squared, and 20 genetic principal components. We fit the linear mixed model on SNP-array-genotyped autosomal variants with MAF $> 10^{-4}$ and missingness $< 0.1$.

#### 7.3 Fine-mapping using approximate conditional analysis

To filter out CNV associations reaching the canonical genome-wide significance threshold ($P < 5 \times 10^{-8}$) that could potentially be explained by linkage disequilibrium (LD) with a more-strongly associated variant within 3Mb—either an imputed SNP or indel [8] or another CNV—we applied a pairwise LD-based filter that we previously developed for identifying likely-causal rare variant associations [11,22].

We implemented the same pairwise LD filter that we used in our previous work by computing the following approximate $\chi^2$ test statistic for variant $i$ (here, a CNV variable) conditioned on variant $j$ (another variant with a stronger association):

$$\chi^2_{i|j} \approx \chi^2_i (1 - r_{ij} \sqrt{\chi^2_j / \chi^2_i})^2.$$

We required that $\chi^2_{i|j} > 29.7168$, i.e., that the approximate conditional test statistic still reach significance ($P < 5 \times 10^{-8}$) for every nearby more-strongly associated variant $j$.

We note that this approximation holds for variants $i$ and $j$ that are weakly correlated (i.e., $r^2_{ij} \ll 1$), which is true when checking whether rare-variant associations are likely to be explained by more-strongly associated common variants. However, after completing analyses, we realized that a better approximation for the conditional $\chi^2$ statistic is:

$$\chi^2_{i|j} \approx \frac{\chi^2_i(1 - r_{ij}\sqrt{\chi^2_j/\chi^2_i})^2}{1 - r^2_{ij}}$$

which accounts for the reduction in variance of the conditional effect size estimate when the variants $i$ and $j$ are highly correlated [23].

Given that the pairwise LD filter that we implemented is usually only mildly conservative (since the estimate of $\chi^2_{i|j}$ that we computed is too small by a factor of $(1 - r^2_{ij})$), we considered the filter to be adequate for identifying likely-causal CNVs. We did also apply the filter to identify likely-causal coding SNP and indel variants associated with systolic and diastolic blood pressure (Fig. 3d), so we checked that accounting for the reduction in variance of conditional effect size estimates had a minimal impact on this analysis (producing only four additional associations that passed the filter, all being common missense variants with weak effect sizes).

#### 7.4 Filtering and annotating fine-mapped associations

##### 7.4.1 Associations with discrete CNV calls

**Filtering somatic CNVs.** We filtered calls intersecting the following regions (in hg19 coordinates) frequently affected by somatic CNVs:

- Immunoglobulin genes
  - *IGK*: chr2:89,000,000–90,274,235
  - *IGH*: chr14:106,032,614–107,288,051
  - *IGL*: chr22:22,380,474–23,265,085
- T-cell receptor genes
  - *TRG*: chr7:38,279,625–38,407,656
  - *TRB*: chr7:141,998,851–142,510,972
  - *TRA*: chr14:22,090,057–23,021,075
  - *TRD*: chr14: 22,891,537–22,935,569
- *DLEU1* / *DLEU2* locus: chr13:50,556,688–51,297,372

**Annotating syndromic CNVs.** We annotated a CNV association as syndromic if it overlapped a previously-curated pathogenic CNV (from the set of 92 pathogenic CNVs curated by ref. [24]). In more detail:

- CNV categories defined based on 100bp bin-overlap or gene-overlap: annotated as syndromic if the location (i.e., start of 100bp bin for bin-level categories; start of gene for gene-level categories), or the median start or end of the CNV calls within the category (lifted to hg19 coordinates), overlapped a pathogenic CNV ($\pm$100kb) of the same type as the category (DEL or DUP).
- Overall pLoF burden: annotated as syndromic if the median start or end of the relevant gene, or the high-confidence pLoF DEL calls contributing to the burden (lifted to hg19 coordinates), overlapped a pathogenic DEL ($\pm$100kb).

##### 7.4.2 Associations with continuous-valued copy-number estimates

In our initial association results, we observed some associations between blood count phenotypes and continuous-valued measurements of WES read alignments (often in telomeric regions) that appeared to reflect technical artifacts rather than copy-number variation. These associations had the property that the association signal was driven by direct read-depth signal from each sample (which is susceptible to confounding from technical effects specific to a sample's exome sequencing) and was greatly reduced when considering only information derived from haplotype neighbors (for which technical effects of sequencing are expected to be independent).

These observations led us to implement filters based on the association strength $\chi^2_{\text{nbr-only}}$ of an alternative version of continuous-valued copy-number estimation (Section 6) in which in the final iteration of the algorithm, we ignored a sample's own WES read-depth data and set its two allelic copy numbers to the means across neighbors of each of its two haplotypes (equivalent to setting $p$=0 in equation (4) in the final iteration). We implemented the following filters based on $\chi^2_{\text{nbr-only}}$ and the original $\chi^2$ statistic from the regular algorithm:

- We required $\chi^2_{\text{nbr-only}} > 20$, i.e., any association truly driven by common copy-altering structural variation should be well-supported by haplotype neighbors.
- We required $\chi^2_{\text{nbr-only}} > \chi^2/2$, i.e., the strength of the association should not drop by more than twofold when using only haplotype neighbors estimate a sample's copy number.

### 8 Follow-up analyses at highlighted loci

#### 8.1 Height-associated pLoF CNVs

##### 8.1.1 Categorization of newly-implicated genes

To determine which genes for which CNVs contributed to pLoF burden associations with height not discoverable in UK Biobank from pLoF SNPs and indels alone, we examined previous large-scale pLoF SNP/indel analyses of height [3, 25] our previous SNP-array based CNV analysis [11], and OMIM entries.

- **Genes previously implicated:**
  - Previous large-scale pLoF SNP/indel analyses reported *CRISPLD1, DTL, HMGA2, IHH*
  - Previous large-scale SNP-array-based CNV analysis reported *DIS3L2* and *UHRF2*
  - OMIM lists autosomal dominant diseases relevant to height for *GH1* (growth hormone deficiency) and *EXT1* (exostoses)
- **Genes newly implicated:**
  - *CCNF, CDK6, CHSY1, PRKG2, TRANK1, UHRF1, USP14*

###### 8.1.2 *CCNF* intragenic exon 3 duplication

Our HMM-based pipeline called a rare duplication spanning chr16:2,432,900–2,433,200 (containing *CCNF* exon 3) in 149 UKB participants that associated with height and blood cell traits. *CCNF* exon 3 is 107bp and was targeted by a single exome capture probe in UKB WES. To confirm and further characterize this SV, we examined UKB WGS data, finding that carriers showed discordant read pairs indicating a tandem duplication (RF read pairs; Supplementary Fig. 2a,b).

**Precision and recall of HMM-based calls.** The 200K UKB WGS data set allowed us to compute the precision and recall of our HMM-based calls of the *CCNF* exon 3 IED as follows:

- For each individual with WGS data ($n = 200,018$), we counted discordant read pairs (RF read pairs) in the region of interest.
- Discordant read pairs cleanly separated *CCNF* exon 3 IED carriers from noncarriers (59 carriers with $\geq 5$ pairs; 0 or 1 pairs for all other individuals).
- Of these 59 carriers:
  - All individuals with a WES-based HMM call were identified via WGS (0 false positives; 55 such carriers).
  - 4 carriers identified via WGS were not called by our WES-based HMM; 1 such carrier had a different exon 3 IED (based on estimated breakpoint locations).

###### 8.1.3 Replication in BioBank Japan

We used SNP-array-based CNV call set we previously generated in BioBank Japan [11] to attempt replication of height associations involving genes we newly implicated. As described previously, association analyses were run on a 179,420-sample uniform-ancestry subset of BioBank Japan and residuals from a model regressing height on age, sex, genotyping arrays (as factors) and 10 PCs as covariates were computed. These residuals were then inverse-normal transformed and used as outcome variables with the independent variable being carrier status of events of interest.

#### 8.2 *RGL3*

##### 8.2.1 Optimized genotyping of exon 6 partial deletion in UKB WES

We expected that the hypertension-associated partial deletion of *RGL3* exon 6 (overlapping only 55bp of coding sequence) was challenging to call from WES read-depth even using our haplotype-informed approach. To further investigate this association and benchmark the quality of our calls of this deletion, we implemented an optimized WES-based genotyping approach that made use of knowledge about the deletion's breakpoints. Specifically, we:

- Counted the number of off-target reads that aligned to chr19:11,407,500–11,408,500 (hg38) (excluding reads flagged as PCR or optical duplicates). This intronic region was not targeted by exome capture and typically does not contain WES read alignments, but the *RGL3* exon 6 partial deletion (which extends $\sim$1kb upstream of exon 6) causes some captured fragments to include sequence from the right flank of the deletion, generating WES read alignments in this region (Supplementary Fig. 2c).
- Computed read depth at the base pairs immediately before and after each breakpoint of the DEL (using `samtools depth`).

We then performed breakpoint-based genotyping as follows (see diagram in Supplementary Fig. 2c):

- Homozygotes for the DEL were called as individuals with 0 WES read alignments counted in the two WES-targeted 100bp bins affected by the DEL (chr19:11,406,800–11,407,000).
- Heterozygous DEL carriers were identified among remaining individuals as those with either of the following types of breakpoint-based evidence:
  - At least 3 off-target reads aligned to chr19:11,407,500–11,408,500.
  - Drop in WES read-depth of $>5$ at the left breakpoint and an increase in read-depth at the right breakpoint.

**Precision and recall of HMM-based calls.** We used the breakpoint-based genotypes (in the subset of our primary analysis set in the 470K WES release) to evaluate the precision and recall of our HMM-based calls of the 1.1kb *RGL3* deletion (which we defined as DEL calls starting at either chr19:11,406,700 or 11,406,800 and ending at chr19:11,407,000, to account for imprecision on read-depth-based calls). This led to an estimate of recall of 7,124 / 8,111 = 87.8% and precision of 7,124 / 7,133 = 99.9%.

| | HMM-based call | No HMM-based call |
|---|---|---|
| Breakpoint-based call of 1.1kb DEL | 7,124 | 987 |
| No breakpoint-based call | 9 | 429,402 |

Further examination of the 9 individuals with an HMM-based call who did not have a breakpoint-based call showed that 6 of the 9 individuals had breakpoint-based evidence that did not quite reach our call thresholds, so for our final genotyping of the 1.1kb DEL, we augmented the breakpoint-based genotypes with HMM-based *RGL3* pLoF calls with at least 1 off-target read (allowing the HMM-based calls to recover these few samples with weaker breakpoint-based evidence).

##### 8.2.2 Comparison to effects on blood pressure of other nonsynonymous variants

We compared the *RGL3* DEL association with systolic and diastolic blood pressure to exome-wide associations of nonsynonymous variants that passed our fine-mapping filter. In more detail, we ran BOLT-LMM linear mixed model association analysis on systolic and diastolic blood pressure, restricting to individuals in the intersection of our primary analysis set and the 470K WES set, to compute associations with WES-based genotype calls as well as genome-wide imputed SNPs and indels (to use for fine-mapping). We then identified significantly-associated likely-causal nonsynonymous variants as follows:

- Restricted to variants within the file `ukb23158_500k_OQFE.annotations.txt.gz` (a list of variant effect annotations provided on the UKB Research Analysis Platform). Among variants in this file, we removed those annoted as synonymous, retaining those annotated as missense(0/5), missense($\geq$1/5), missense(5/5), or LoF.
- Removed variants with associations potentially explainable by LD to `imp_v3` SNPs or indels based on approximate conditional analysis (see Section 7.3).

##### 8.2.3 Replication of hypertension association in *All of Us*

We genotyped the *RGL3* 1.1 kb deletion in *All of Us* participants with available WGS data by detecting chimeric sequence created by the deletion, identifying reads that align to one flank of the deletion and contain sequence from the other flank (juxtaposed by the deletion). Specifically, we:

- Extracted short-read WGS alignments overlapping either chr19:11,406,805–11,406,809 or chr19:11,407,947-11,407,951
- Restricted to reads with SAM flags 99, 147, 83, 163, 97, 145, 81, or 161
- Counted the number of reads containing the chimeric sequence "CTTCCAAGTG"

To help distinguish individuals homozygous for the deletion from heterozygous carriers, we also counted the number of reads (with the same SAM flags as above) mapping to chr19:11,406,809–

11,407,946 (within the 1.1kb deletion), normalizing this read count by the number of reads mapping to chr19:11,355,386-11,365,698 (the nearby *PLPPR2* gene, which did not contain common CNVs in gnomAD-SV [17]).

We found that the deletion had an overall allele frequency of 0.0045 in *All of Us* (as it was much rarer in non-European ancestries). Chimeric reads cleanly identified carriers and non-carriers of the deletion (Supplementary Fig. 2d):

| | | Number of chimeric reads | | | | $\frac{\text{Read count in 1.1kb DEL}}{\text{Read count in nearby 10kb region}}$ | |
|---|---|---|---|---|---|---|---|
| Deletion | $n$ | mean | s.e.m. | min | max | mean | s.e.m. |
| Non-carrier | 243,211 | $7.4 \times 10^{-5}$ | $1.74 \times 10^{-5}$ | 0 | 1 | 0.109 | $1.8 \times 10^{-5}$ |
| Carrier | 2,183 | 19.3 | 0.117 | 3 | >40 | 0.0543 | 0.000159 |

We then tested the deletion for association with hypertension status in *All of Us* (defined as presence of at least one EHR-derived record of hypertension) adjusting for:

- `sex_at_birth` found within `genomic_metrics.tsv`
- `ancestry_pred` and 16 PCs found within `ancestry_preds.tsv`
- age (based on birth year)

##### 8.2.4 Effect of the *RGL3* exon 6 partial deletion on *RGL3* expression in GTEx

We identified 7 carriers of the *RGL3* 1.1kb deletion in GTEx v8 using the same approach as above to search for chimeric sequence created by the deletion (Section 8.2.3).

Across the 45 tissues with at least 1 carrier, we found that the majority of normalized effect sizes of the *RGL3* deletion on *RGL3* expression were negative (36/45 negative; binomial test $P = 6.6 \times 10^{-5}$). This seems to suggest that the apparent LoF effect of the *RGL3* 1.1kb deletion may be mediated in part by reduced expression of *RGL3* alleles carrying the deletion; however, further work will be required to fully elucidate the mechanism.

##### 8.2.5 Association tests of *RGL3* DEL with binary phenotypes

The protective effect of the *RGL3* 1.1kb deletion on hypertension, together with the observation of 37 UKB participants homozygous for the deletion, suggested the possibility that *RGL3* (or a pathway in which it functions) could be a potential drug target. To further explore this possibility, we checked whether *RGL3* deletion status associated with any adverse effects on health. Specifically, we ran association tests against the 1,129 "first-occurrence" disease phenotypes curated by UK Biobank using the BinomiRare test [26] (using the same analytical setup described in Section 10). For each disease phenotype, we ran both a test on only *RGL3* DEL homozygote status (i.e., 37 homozygotes coded as 1, all other individuals coded as 0) and a test on *RGL3* DEL carrier/noncarrier status.

*RGL3* DEL homozygotes did not exhibit any Bonferroni-significant associations with diseases (minimum $P$=0.001), suggesting that the 37 homozygous UKB participants were generally healthy (though this test did not have power to detect subtle effects on disease risk). Similarly, *RGL3* DEL carriers did not have any Bonferroni-significant associations with increased risk of any disease. We did observe one marginally Bonferroni-significant association with reduced risk of the ICD-10 category F17 (mental and behavioral disorders due to use of tobacco) ($P = 3.2 \times 10^{-5}$, OR=0.87 (0.83–0.91)) along with the protective association with hypertension. The *RGL3* DEL did not associate with smoking status, such that the association with reduced hypertension risk was independent of the possible association with reduced F17 risk.

#### 8.3 *CTRB2*

##### 8.3.1 Validation of UK Biobank WES-based genotyping using WGS

We confirmed the validity of our WES read-depth-based genotypes of the *CTRB2* exon 6 deletion by comparing WES-derived copy-number estimates to normalized WGS read-depth for 500 UK Biobank participants (Fig. 4h). We used mosdepth [15] to measure WGS read-depth across chr16:75,204,723–75,205,296 and chr16:75,223,665–75,224,238 (the latter being the homologous region in *CTRB1*), summed these values, and normalized by genome-wide WGS coverage.

##### 8.3.2 Replication of T2D association in *All of Us*

We genotyped the *CTRB2* exon 6 deletion in *All of Us* by counting the number of reads with chimeric sequence supporting the deletion. In more detail, we:

- Extracted short-read WGS alignments overlapping either chr16:75,204,710–75,204,725 or chr16:75,205,290–75,205,310
- Restricted to reads with SAM flags 99, 147, 83, 163, 97, 145, 81, or 161
- Counted the number of reads containing the chimeric sequence "AAAGCCCAGACCCCA"

To help distinguish individuals homozygous for the deletion from heterozygous carriers, we also counted the number of reads (with the same SAM flags as above) mapping to chr16:75,204,725–75,205,275 (within the deletion). We normalized both read counts by each individual's genome-wide mean WGS coverage (`mean_coverage` found within `genomic_metrics.tsv`). These normalized counts of chimeric reads and within-deletion reads allowed us to cleanly genotype the *CTRB2* deletion (Supplementary Fig. 2e).

We then tested the deletion for association with type 2 diabetes status in *All of Us* (defined as presence of at least one EHR-derived record of T2D) adjusting again for:

- `sex_at_birth` found within `genomic_metrics.tsv`

- `ancestry_pred` and 16 PCs found within `ancestry_preds.tsv`
- age (based on birth year)

#### 8.4 *FCGR3B*

##### 8.4.1 Optimized genotyping of *FCGR3A* / *FCGR3B* copy number in UKB WES

Based on previous literature [27], most *FCGR3A* CNVs also contain *HSPA6*, and most *FCGR3B* CNVs also contain *HSPA7*. We therefore optimized our genotyping of *FCGR3A* and *FCGR3B* copy number from UKB WES data by computing normalized WES read counts in the following regions (restricting to alignments with positive mapping quality):

- *HSPA6+FCGR3A*: chr1:161,524,000-161,550,500
- *HSPA7+FCGR3B*: chr1:161,606,000-161,632,000

We then refined the normalized read counts using our phasing pipeline and estimated copy-number of *FCGR3A* and *FCGR3B* based on modes of these distributions, which were well-separated (Fig. 5c).

##### 8.4.2 Validation of *FCGR3B* genotypes using UKB proteomic and WGS data

We validated our genotyping of *FCGR3B* copy number by examining our WES-based copy-number estimates for consistency with FCGR3B protein abundances recently measured in plasma [28] (Supplementary Fig. 5). We restricted analysis to individuals in our primary analysis set with protein levels available ($n$=49,364). We regressed Normalized Protein eXpression (NPX) values for FCGR3B on *FCGR3B* copy number and controlled for age, sex, and 20 PCs (as NPX was already normalized for technical covariates). We then converted all estimates to the linear scale through the transformation $2^{\text{NPX}}$.

We further validated our WES-based genotyping by comparing WES-derived copy-number estimates to normalized WGS read-depth for 500 UK Biobank participants (Supplementary Fig. 5). We counted WGS reads aligning to *HSPA7+FCGR3B* (chr1:161,606,000-161,632,000) with SAM flags 99, 147, 83, 163, 97, 145, 81, or 161 and positive mapping quality, and we normalized by genome-wide WGS coverage.

#### 8.5 *SIGLEC14–SIGLEC5* gene fusion

##### 8.5.1 Genotyping *SIGLEC14–SIGLEC5* CNVs in UK Biobank WES

Our common CNV analysis pipeline (Section 6) had already generated continuous-valued copy-number measurements for the region chr19:51,630,200–51,646,900 (based on a known CNV), and these measurements (based on counting WES read alignments with any mapping quality) exhibited clear separation of modes corresponding to copy-number states. Genotyping based on these modes led to allele frequencies of 0.17 and 0.01 for the *SIGLEC14–SIGLEC5* fusion (deletion) and reciprocal duplication, respectively.

##### 8.5.2 Genotyping and gene expression analysis in GTEx

To identify carriers of the *SIGLEC14-SIGLEC5* gene fusion in GTEx v8, we extracted read counts from the deletion region (chr19:51,630,200–51,646,900) as well as within a control gene (*ZNF175*; chr19:51,571,283-51,592,510), restricting to reads with SAM flags 99, 147, 83, 163, 97, 145, 81 or 161. Normalizing the within-deletion read count against the control-gene read count produced well-separated copy-number modes, as expected given the large size of the CNV ($\sim$17kb).

Among 879 GTEx donors with WGS data, we identified 38 individuals homozygous for the gene fusion, 258 individuals heterozygous for the gene fusion, 565 individuals with no CNV (i.e., hom-REF), and 18 individuals with the reciprocal duplication, leading to allele frequencies of 0.19 and 0.01 for the fusion and duplication, respectively.

For each tissue in GTEx, we then:

- Extracted TPM and normalized expression values for *SIGLEC14* and *SIGLEC5* as well as standard GTEx eQTL covariates
- Computed the median ratio of *SIGLEC14* expression to *SIGLEC5* expression (using TPM values) among individuals with no CNV (hom-REF)
- If the tissue had normalized expression values available for both *SIGLEC14* and *SIGLEC5*:
  - Removed individuals with *SIGLEC14* duplication
  - Computed allelic fold change estimates [29] controlling for covariates. In more detail:
    - Regressed *SIGLEC14* (resp., *SIGLEC5*) TPM on fusion genotype (0/1/2) using standard GTEx covariates (mean-centered) and used the estimated covariate coefficients to compute covariate-adjusted TPM values for each individual. We then used bootstrap to estimate the confidence interval for allelic fold change (re-estimating the intercept and the coefficient of the fusion genotype in each bootstrap sample, and using these values to compute allelic fold change [29]).
  - Normalized effect size estimates: Regressed normalized *SIGLEC14* (resp., *SIGLEC5*) expression on fusion genotype (0/1/2) controlling for covariates.

### 9 Analysis of paralogous sequence variants at 7q22.1 and *DEFA1* / *DEFA3*

#### 9.1 Genotyping paralogous sequence variants (PSVs) from WGS read alignments

SNP and indel variants within segmental duplication regions (i.e., PSVs) are difficult to genotype from short-read sequencing data for two main reasons: (i) reads may align non-uniquely to multiple paralogous regions, such that accurate analysis requires collating alignments at all paralogous bases and analyzing them together; and (ii) copy number of duplicated segments can be high and can vary across individuals, such that many more genotypes than the usual three (AA/AB/BB) are possible, and short-read WGS/WES data from a single sample does usually provide enough information to distinguish the possible genotypes.

To overcome these challenges, we employed an approach in which we first roughly estimated PSV genotypes from WGS read alignments (available for ∼200K UKB participants) and then used our haplotype-informed approach to denoise PSV genotypes and impute them into the remainder of the UKB cohort.

**Estimating PSV genotypes from WGS read alignments.** We applied the following approach (depicted in Supplementary Fig. 4) to estimate PSV genotypes from short-read WGS data:

1. Identify all regions in the GRCh38 reference genome with high sequence similarity to the duplicated segment of interest. Such regions effectively act as ”baits” for reads: i.e., reads originating from our region of interest may map to these other, similar, regions.
    - To detect these bait regions, we first divided the region of interest into 5kb blocks.
    - For each block, we identified highly similar regions across the genome using BLAT [30] (`-minScore=200 -minIdentity=98`).
    - We manually reviewed the BLAT output to select genomic segments to include in the final bait set and to determine the number of bait segments aligned to each position in the duplicated segment of interest. (This number could vary across the segment, e.g., if part of the segment had a paralog elsewhere in the genome, or if alleles start or end in the middle of a segment.)
2. Extract all reads originally mapped to any of the bait regions; then realign these reads to a reference sequence consisting of only one copy of the duplicated segment (plus a small buffer sequence containing the beginning of a second copy of the same segment). Realigning reads in this way circumvented complexities arising from sequence differences between copies of

the segment in the reference that could could interfere with harmonization of the original alignments to GRCh38. We extracted previously-mapped reads using samtools [4] and we realigned these reads using bwa [31].

3. Ascertain and estimate PSV allele fractions for all common sequence variants in the segment.

   - For each sample, we generated "pileups" counting the numbers of reads supporting sequence variants (i.e., alternate bases or indels) observed among the realigned reads (using `htsbox -Q20 -l50`). We considered a sample to be a likely carrier of a PSV if it was supported by at least 5 alternate reads. We then collated across samples and identified common PSVs as those with carrier frequency >0.02.
   - For each sample, for each common PSV identified above, we estimated its PSV allele fraction (PSVAF; i.e., the fraction of repeat units containing the PSV) as the fraction of reads in the pileup that supported the PSV allele.

4. Convert PSV allele fractions to absolute copy number estimates. For each sample, for each PSV, we estimated PSV copy number (PSVCN; i.e., the number of repeat units containing the PSV) by multiplying PSVAF by the individual's estiamted total copy number at the PSV position. We obtained estimates of total segmental copy number from our normalized WES read count analysis pipeline (Section 6) as it produced well-separated, evenly spaced copy-number modes when applied to 7q22.1 (chr7:102340000-102691000) and *DEFA1 / DEFA1B / DEFA3* (chr8:6977200-6979400 / chr8:6996300-6998500 / chr8:7015400-7017600). We applied position-specific "bonus CN" adjustments to total copy number to account for differences in the numbers of bait regions in GRCh38 paralogous to different portions of the segment.

**Denoising and imputing PSV genotypes.** For each common PSV at which we estimated PSVCN (continuous-valued) using the above procedure, we denoised PSVCN estimates using the phasing algorithm described in Sections 6.2–6.3. This haplotype-sharing analysis also allowed us to simultaneously impute PSVCN estimates into the remainder of the UKB cohort (i.e., participants not in the 200K WGS data set) by setting each target haplotype's (haploid) PSVCN to the mean of its haplotype neighbors.

#### 9.2 7q22.1 segmental duplication

##### 9.2.1 Association and fine-mapping with T2D and chronotype in UKB

**UK Biobank phenotype refinement.** Copy-number measurements in the 7q22.1 segmental duplication region from our common-CNV pipeline associated with HbA1c, leading us to perform follow-up analyses of association with type 2 diabetes status in UK Biobank. We also noticed that the lead SNP at the locus had previously also been associated with chronotype [32], leading us to examine this phenotype as well. We coded T2D status and chronotype as follows:

- Type 2 diabetes: We identified likely cases of T2D as individuals who had reported diabetes diagnosed by a doctor (Data-Field 2443) and did not report type 1 diabetes, gestational diabetes, or diabetes insipidus (Data-Field 20002). Individuals who reported doctor diagnosed diabetes and also reported type 1 diabetes, gestational diabetes, or diabetes insipidus were removed from the analysis.

- Chronotype: We used Data-Field 1180 to define chronotype, coding "Definitely a 'morning' person" as 1, "More a 'morning' than 'evening' person" as 2, "More an 'evening' than a 'morning' person" as 3, and "Definitely an 'evening' person" as 4. Individuals who selected "Do not know" or "Prefer not to answer" were removed from the analysis.

**Association and fine-mapping analyses including PSVs.** We applied the PSV analysis pipeline described above to estimate copy numbers of PSVs within the 7q22.1 segmental duplication region. We then tested PSVs for association with T2D and chronotype using linear regression implemented in BOLT-LMM, restricting to the same samples and including the same covariates as in our primary association analysis pipeline (for 57 quantitative traits).

To facilitate fine-mapping, we also computed linear regression association statistics for imputed SNPs and indels in the region, including genotyping array, WES release, sex, age, age squared, and 20 PCs as covariates, and applying standard MAF and INFO filters (`--bgenMinMAF=1e-3` and `--bgenMinINFO=0.3`).

For many PSVs, association strengths of PSV copy numbers considerably exceeded those of the overall copy number of the 99kb segment as well as imputed SNPs and indels flanking the duplication region (Fig. 4c). To identify potential causal variants, we performed fine-mapping using SuSiE [33] on PSVs together with SNPs and indels (restricting to significantly-associted variants for computational efficiency, and running SuSiE on phenotypes and genotypes residualized for our standard covariate set).

Using SuSiE to fine-map T2D associations, we observed that regardless of the number of causal variants allowed in the model (ranging from 1 to 5 variants), SuSiE consistently chose only 1 variant (the *RASA4* Y731C missense PSV; chr7:102,485,072 T>C in GRCh38 coordinates) with PIP = 0.99. Fine-mapping results for chronotype were less robust (as the *RASA4* Y731C missense PSV and other PSVs within segmental duplication exhibited similar association strengths; Fig. 4c), and SuSiE's selection of variants varied somewhat depending on modeling parameters.

##### 9.2.2 Replication of T2D association in *All of Us*

We measured copy number of the 99kb segmental duplication as well as the *RASA4* missense PSV in *All of Us* as follows:

- Extracted short-read WGS alignments overlapping either chr7:102,341,679–102,357,966 or chr7:102,473,938–102,691,171
- Restricted to reads with SAM flags 99, 147, 83, 163, 97, 145, 81, or 161 and computed the read count
- Used `htsbox` to count reads supporting the C allele at chr7:102,485,072, chr7:102,584,251, and chr7:102,683,312 (locations of the PSV in the segmental duplications present in GRCh38; T is the reference allele)

Normalizing both of the above read counts by each individual's genome-wide mean WGS coverage (`mean_coverage` found within `genomic_metrics.tsv`) produced estimates of copy number of the 99kb segment and the PSV (up to scaling constants) that exhibited a multimodal distribution as expected. We then tested these continuous-valued estimates of 99kb segment copy number and PSV copy number for association with type 2 diabetes status in *All of Us* using the same analytical setup as for *CTRB2* (Section 8.3.2).

#### 9.3 *DEFA1 / DEFA3* segmental duplication

We applied the same PSV genotyping, association, and fine-mapping analysis as above at the *DEFA1 / DEFA3* segmental duplication locus, at which our common-CNV association screen had detected associations with basophil count and monocyte count. Like at 7q22.1, PSV association strengths with both phenotypes considerably exceeded those of the overall copy number of the 19kb repeat as well as imputed SNPs and indels flanking the repeat region (Fig. 4e).

Using SuSiE to fine-map the basophil count and monocyte count associations, we observed that regardless of the number of causal variants allowed in the model (ranging from 1 to 5 variants), the first selected cluster of variants was always a subset of PSVs from a 5-SNP haplotype in an Alu element within the 19kb repeat. The five PSVs and their LD are listed below; base pairs are in GRCh38 chr8 coordinates (corresponding to one particular repeat unit in the GRCh38 assembly).

| LD ($r$) | 6,993,547:C/A | 6,993,555:G/A | 6,993,595:C/T | 6,993,602:C/T | 6,993,671:G/A |
|---|---|---|---|---|---|
| 6,993,547:C/A | 1 | 0.998 | 0.989 | 0.989 | 0.975 |
| 6,993,555:G/A | | 1 | 0.990 | 0.990 | 0.976 |
| 6,993,595:C/T | | | 1 | 0.998 | 0.982 |
| 6,993,602:C/T | | | | 1 | 0.984 |
| 6,993,671:G/A | | | | | 1 |

### 10 Association tests with disease phenotypes

We ran gene-level burden analyses that collapsed all types of pLoF variants (CNVs, SNPs, and indels) to maximize power to detect rare loss-of-function effects. We ran association tests with binary traits using the BinomiRare test [26] to obtain $P$-values robust to case-control imbalance while adjusting for age, sex, and oligo batch. As previously described [11, 34], for computational efficiency, we reimplemented the BinomiRare test and applied a binomial approximation when the number of observed cases among carriers exceeded 100. We restricted disease association analyses to our primary European-ancestry sample set that passed quality control filters.

We tested two sets of binary traits: 1,129 "first-occurrence" binary disease phenotypes curated by UK Biobank, and 11,250 ICD-10 codes (3-character and 4-character prefixes of reported ICD-10 codes) reported in Hospital Episode Statistics (HES) or cancer or death registries.

We examined associations that reached $P < 5 \times 10^{-8}$ and were not driven by syndromic or somatic CNVs (Section 7.4.1). These associations included 68 pLoF gene-trait associations that did not reach the significance threshold when analyses were restricted to pLoF SNPs and indels alone, of which nearly all were known gene-trait associations, mostly corresponding to Mendelian disorders (Supplementary Data 5). The remaining few associations were near the (loose) significance threshold, so we did not investigate further; larger cohorts or case-control studies will be needed to power additional discovery.

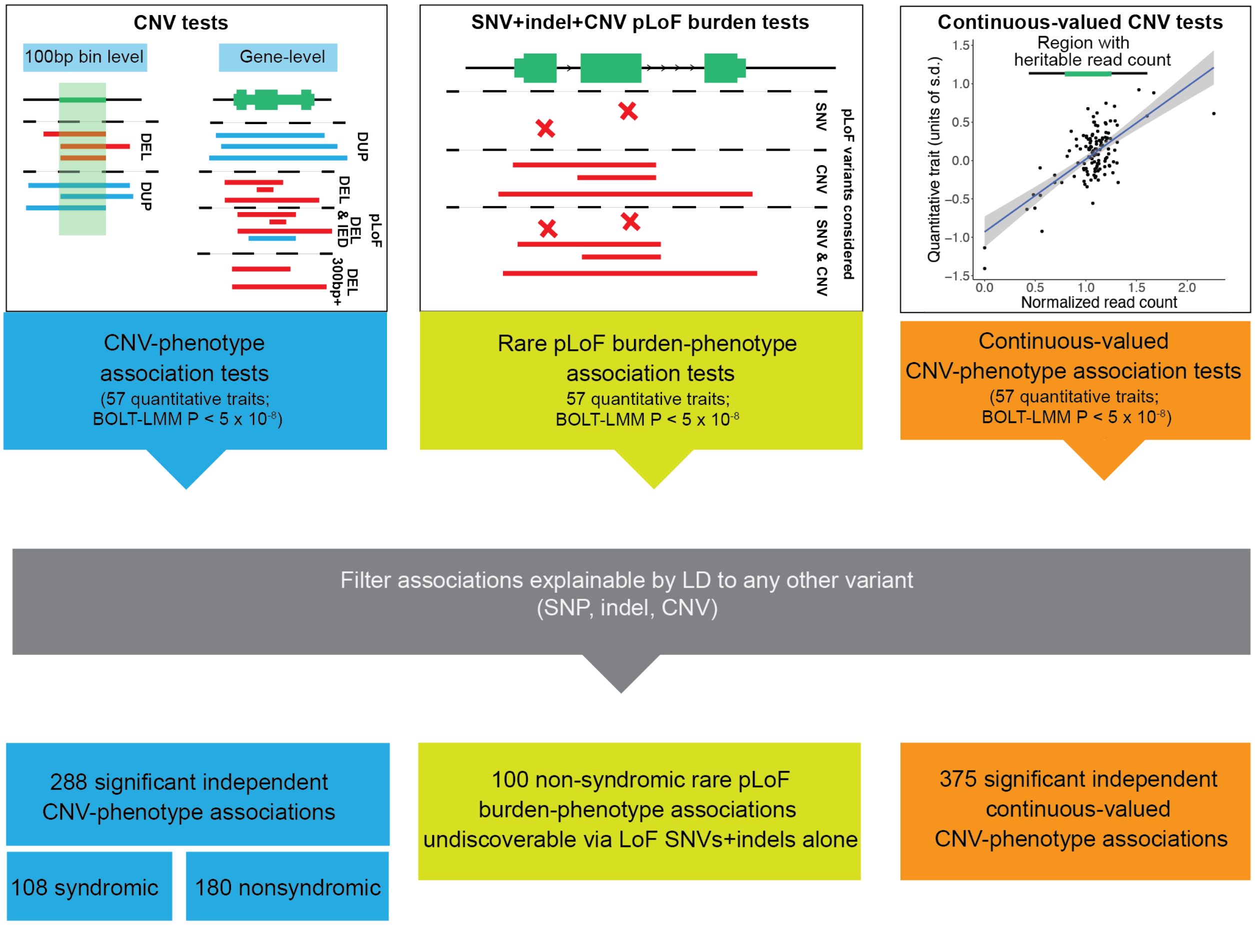

**Supplementary Figure 1. Overview of primary association analyses of 57 heritable quantitative traits.** Categories of variants and CNV measurements tested are depicted, and summary numbers of results from each set of association tests that remained after filtering are provided at the bottom.

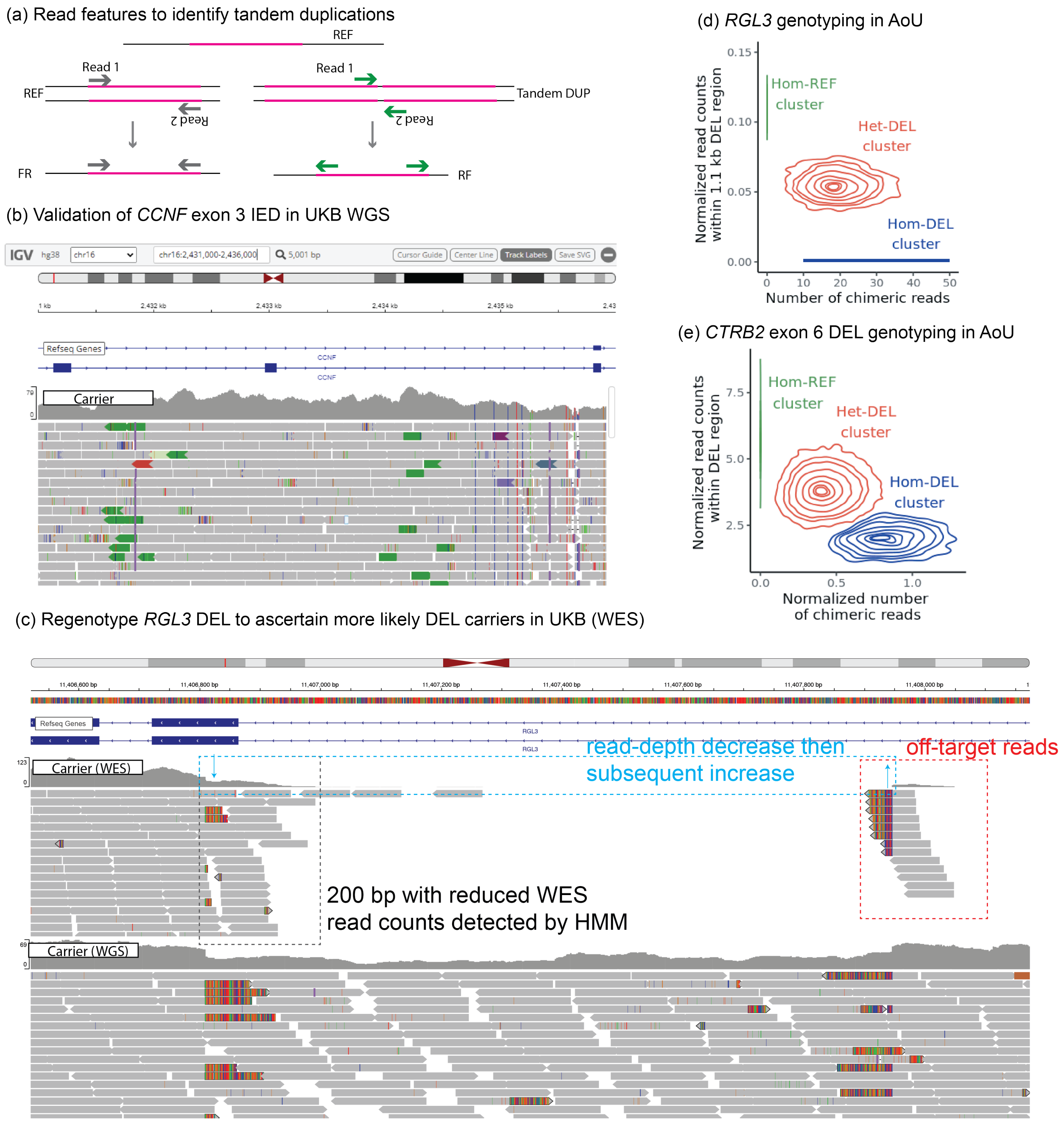

**Supplementary Figure 2. Additional CNV genotyping at key loci.** (a) Schematic of discordant WGS reads that confirm tandem duplications and indicate breakpoint locations. (b) We genotyped the *CCNF* exon 3 IED using discordant WGS reads (shown for a carrier in UKB) to assess precision and recall of WES-based calls from our HMM. (c) IGV tracks of WES and WGS alignments for an *RGL3* deletion carrier. Top: WES features used in optimized breakpoint-based genotyping; bottom: independent confirmation of deletion from WGS. (d, e) In *All of Us* (AoU), chimeric WGS reads and within-deletion read counts allowed the *RGL3* and *CTRB2* deletions to be cleanly genotyped. (The homozygous deletion cluster for *RGL3* contained $<20$ carriers, so to comply with AoU policy, the Hom-DEL line depicted is predicted from the heterozygous cluster.)

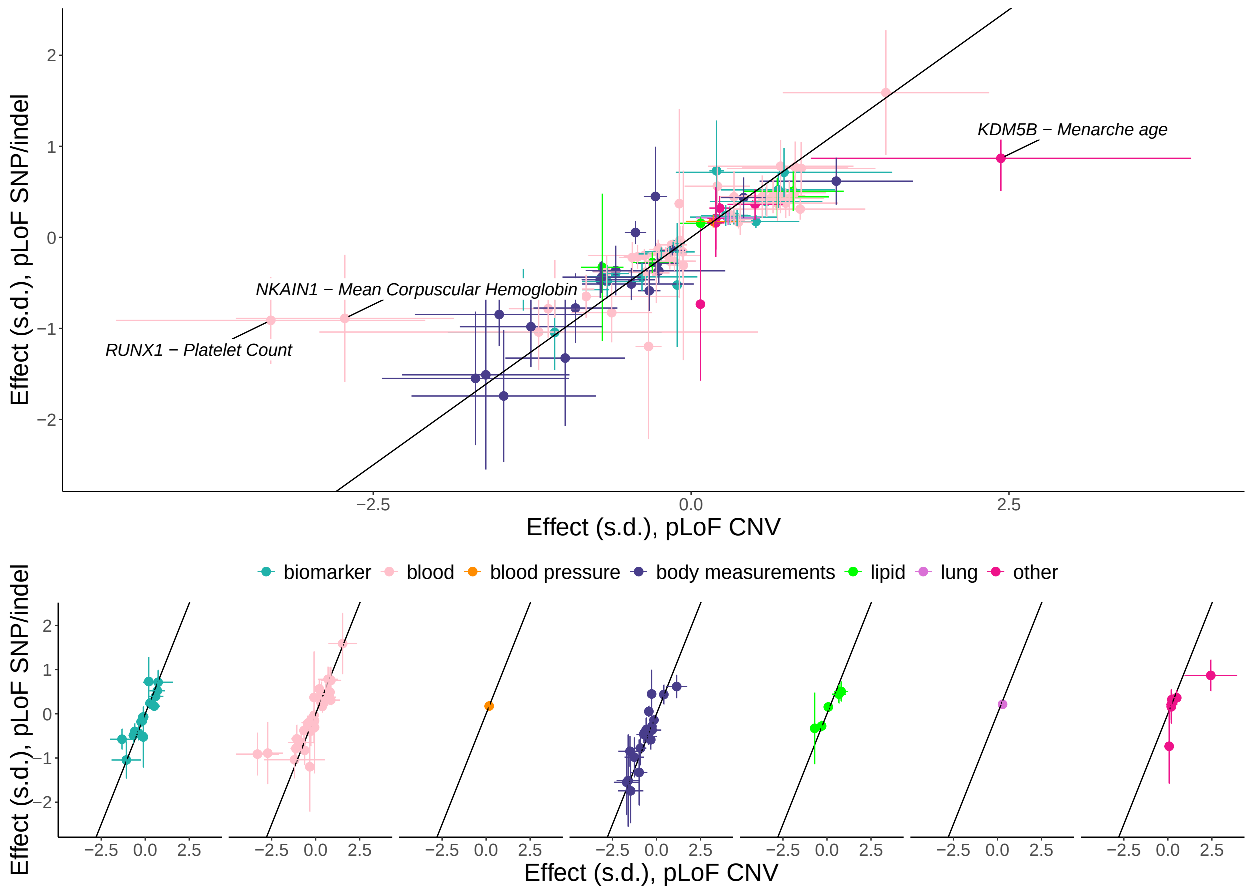

**Supplementary Figure 3. Consistency of effect sizes of pLoF CNV and SNP/indel variants across gene-trait associations.** Data are shown for all associations discovered only upon considering pLoF CNVs (i.e, not reaching significance in SNP/indel-only burden tests). The top plot is a merge across all traits, and the bottom plots show each phenotype category separately. Error bars are 95% confidence intervals.

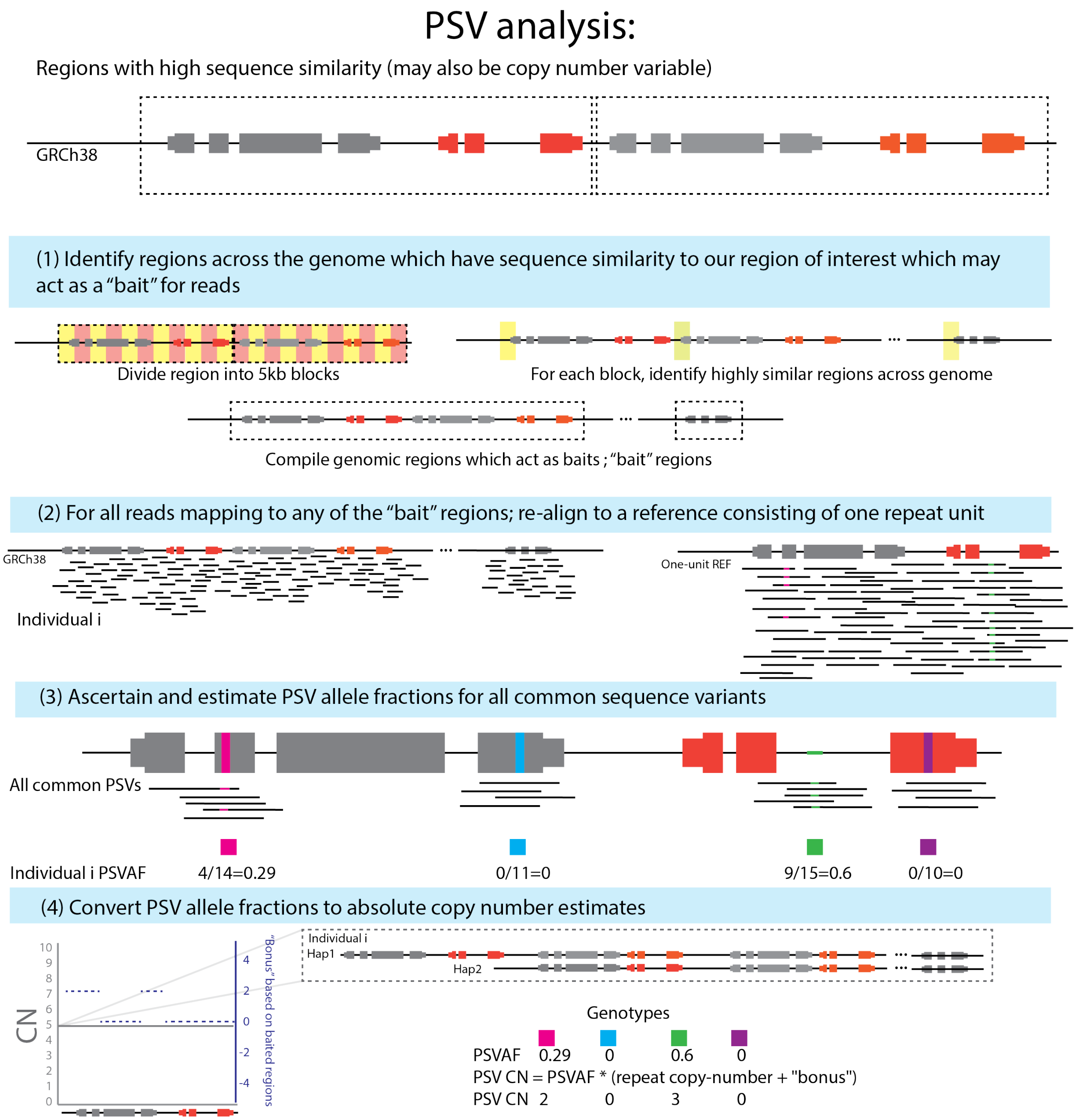

**Supplementary Figure 4. Overview of copy-number estimation for paralogous sequence variants (PSVs).** This figure provides a graphical overview of the pipeline we used to estimate copy-numbers of PSVs—i.e., SNPs and indels carried on one or more copies of a multi-copy segment—from WGS read alignments (Section 9). We then refined these estimates using haplotype-sharing information.

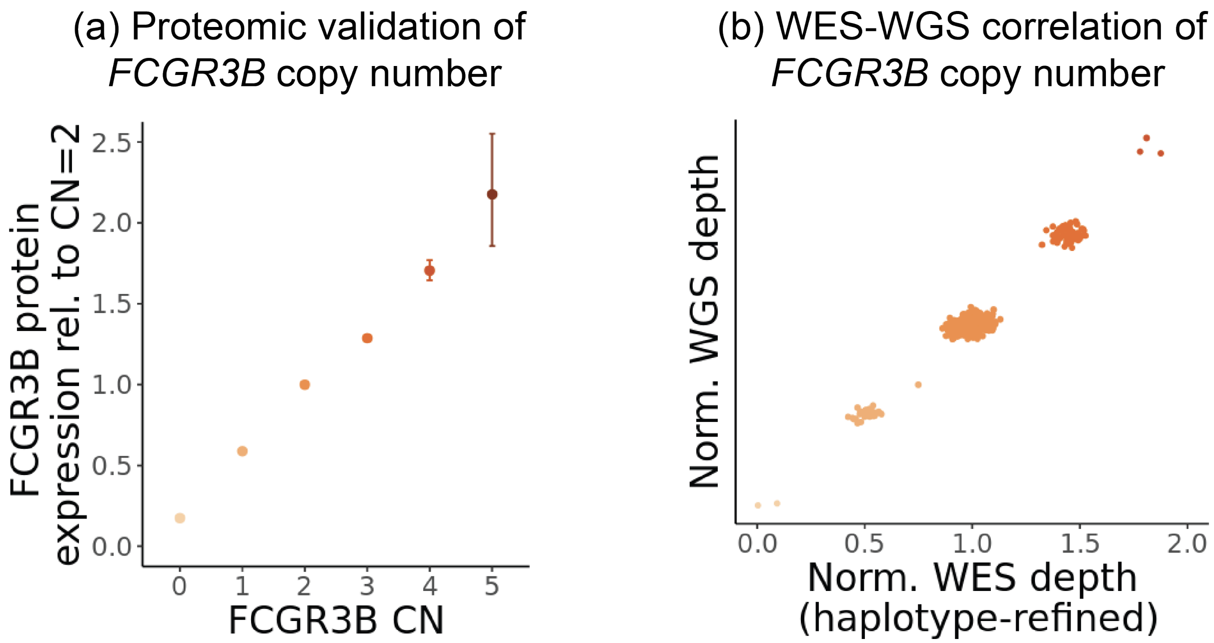

**Supplementary Figure 5. Validation of *FCGR3B* copy number estimation.** (a) Normalized protein expression of FCGR3B for each *FCGR3B* copy-number state relative to CN=2. Estimates and 95% confidence intervals were obtained from linear regression analyses of NPX values and then converted to the linear scale ($2^{NPX}$). (b) Scatter plot of normalized whole-genome and whole-exome sequencing read depths at *FCGR3B* for 500 UKB participants.

| | # DEL | # DUP |
|---|---|---|
| All | 65.75 | 27.69 |
| Size (bp) | | |
| ≤500bp | 45.03 | 6.76 |
| 500bp-5kb | 8.93 | 9.05 |
| >5kb | 11.78 | 11.88 |
| Size (100bp bins) | | |
| 1 | 28.51 | 0.00 |
| 2 | 9.53 | 3.06 |
| 3 | 6.11 | 3.57 |
| 4 | 3.30 | 2.58 |
| 5+ | 18.29 | 18.48 |
| Number of genes | | |
| 0 | 33.57 | 5.33 |
| 1 | 27.45 | 18.16 |
| 2 | 3.07 | 2.57 |
| 3 | 1.52 | 1.18 |
| 4 | 1.05 | 1.28 |
| 5+ | 1.18 | 1.38 |
| Number of exons | | |
| 0 | 33.57 | 5.33 |
| 1 | 18.75 | 8.84 |
| 2 | 5.17 | 3.40 |
| 3 | 1.78 | 1.94 |
| 4 | 1.28 | 1.52 |
| 5 | 1.21 | 1.33 |
| 6 | 1.78 | 1.37 |
| 7 | 1.39 | 1.15 |
| 8 | 1.47 | 1.12 |
| 9 | 1.07 | 1.53 |
| 10+ | 2.04 | 4.09 |

**Supplementary Table 1. Number of HMM-based CNV calls per individual.** Average number of calls per individual made in UK Biobank for deletions (DEL) and duplications (DUP), as a function of size (in both bp and number of 100bp bins used) as well as number of genes (or exons) overlapped by calls.

| | DEL | DUP |
|---|---|---|
| All | 0.89 | 0.91 |
| Size (bp) | | |
| ≤500bp | 0.88 | 0.86 |
| 500bp-5kb | 0.92 | 0.92 |
| >5kb | 0.90 | 0.95 |
| Size (100bp bins) | | |
| 1 | 0.84 | – |
| 2 | 0.89 | 0.74 |
| 3 | 0.92 | 0.88 |
| 4 | 0.92 | 0.93 |
| 5+ | 0.95 | 0.95 |

**Supplementary Table 2. Validation of HMM-based CNV calls based on consistency in WGS read-depth.** HMM-based CNV calls (made from WES read counts) were examined for consistent read-depth deviation in WGS of 100 UK Biobank participants (i.e., reduced read-depth for deletions and increased read-depth for duplications). Validation rate was estimated as the difference between the proportions of calls with consistent vs. inconsistent WGS read-depth (Section 5.4). Estimated validation rates are shown for deletions (DEL) and duplications (DUP) as a function of size (in both bp and number of 100bp bins used).

| Trait | Model | OR (95% CI) or $\beta$ (s.e.) | $P$ |
|---|---|---|---|
| Hypertension | Logistic | 0.86 (0.82–0.90) | $6.3 \times 10^{-10}$ |
| Systolic BP | BOLT-LMM | -0.11 (0.01) | $6.1 \times 10^{-23}$ |
| Diastolic BP | BOLT-LMM | -0.11 (0.01) | $3.9 \times 10^{-25}$ |

**Supplementary Table 3. Associations of *RGL3* exon 6 partial deletion with hypertension and blood pressure.** These data correspond to the refined breakpoint-based genotypes of the *RGL3* 1.1kb deletion (Section 8.2) (in contrast to the association data reported in Supplementary Data 2 for read-count-based CNV calls). All models controlled for WES release, assessment center, genotype array, sex, age, age squared, and 20 PCs as covariates. OR, odds ratio.

**(a) *RGL3* 1.1kb DEL association with hypertension**

| | Sample size ($n$) | | | | | | |
|---|---|---|---|---|---|---|---|
| Ancestry | Total | Cases | Controls | AF | OR | 95% CI | $P$ |
| All | 245394 | 82180 | 163214 | 0.00447 | 0.833 | (0.7548–0.9186) | 0.000259 |
| eur | 133581 | 48288 | 85293 | 0.00703 | 0.818 | (0.7362–0.9095) | 0.0002 |
| afr | 56913 | 20366 | 36547 | 0.00119 | 0.895 | (0.608–1.318) | 0.575 |
| amr | 45035 | 11807 | 33228 | 0.00178 | 1.02 | (0.6916–1.498) | 0.929 |

**(b) 7q22.1 associations with T2D**

| | Sample size ($n$) | | | 99kb segment CN | | *RASA4* Y731C PSV CN | |
|---|---|---|---|---|---|---|---|
| Ancestry | Total | Cases | Controls | $P$ | sign | $P$ | sign |
| All | 245394 | 36797 | 208597 | 0.000201 | + | $2.84 \times 10^{-5}$ | + |
| eur | 133581 | 17955 | 115626 | 0.00699 | + | 0.000999 | + |
| afr | 56913 | 10277 | 46636 | 0.153 | + | 0.107 | + |
| eas | 5706 | 461 | 5245 | 0.319 | - | 0.628 | - |
| amr | 45035 | 7631 | 37404 | 0.00507 | + | 0.00901 | + |
| mid | 942 | 129 | 813 | 0.426 | - | 0.153 | - |
| sas | 3217 | 344 | 2873 | 0.45 | + | 0.835 | + |

**(c) *CTRB2* exon 6 DEL association with T2D**

| | Sample size ($n$) | | | | | | |
|---|---|---|---|---|---|---|---|
| Ancestry | Total | Cases | Controls | AF | OR | 95% CI | $P$ |
| All | 245394 | 36797 | 208597 | 0.0603 | 0.926 | (0.8936–0.9595) | $2.26 \times 10^{-5}$ |
| eur | 133581 | 17955 | 115626 | 0.0777 | 0.933 | (0.8935–0.9746) | 0.0018 |
| afr | 56913 | 10277 | 46636 | 0.0229 | 0.928 | (0.8322–1.036) | 0.183 |
| eas | 5706 | 461 | 5245 | 0.029 | 0.614 | (0.3691–1.023) | 0.061 |
| amr | 45035 | 7631 | 37404 | 0.0614 | 0.909 | (0.8398–0.9834) | 0.0175 |
| mid | 942 | 129 | 813 | 0.0377 | 1.36 | (0.6623–2.806) | 0.4 |
| sas | 3217 | 344 | 2873 | 0.0446 | 1.06 | (0.6989–1.604) | 0.787 |

**Supplementary Table 4. Replication of associations with hypertension and T2D in *All of Us*.** Analyses that combined samples across ancestries ("All"; top row of each subtable) controlled for sex at birth, predicted ancestry category, age (based on year of birth), and 16 genetic PCs; ancestry-specific analyses controlled for the same covariates excluding predicted ancestry. The association of the *RGL3* 1.1kb deletion with hypertension and the *CTRB2* exon 6 deletion with T2D were computed using a logistic model, whereas the associations of the 7q22.1 99kb segment duplication copy number and the *RASA4* Y731C PSV copy number were computed using a linear model.

| Autosomal genes | All | No coding exon overlapped | Coding exon overlapped | Entire gene covered |
|---|---|---|---|---|
| | $n$ (proportion) | | | |
| All | 18,651 | 16,546 (88.71%) | 2,105 (11.29%) | 319 (1.71%) |
| With non-NA pLI score | 16,586 | 14,859 (89.59%) | 1,727 (10.41%) | 207 (1.25%) |
| | Mean pLI score (s.e.m.) | | | |
| With non-NA pLI score | 0.24 (0.003) | 0.25 (0.0032) | 0.16 (0.0077) | 0.09 (0.014) |

**Supplementary Table 5. Proportion of genes affected by common copy-altering structural variation.** Number and proportion of autosomal genes with no coding exon, at least one coding exon, or the entire gene overlapped by a region we identified as likely to harbor common copy-altering variants and subsequently analyzed (Section 6). Mean pLI score, and standard error of the mean, are reported across genes in each category (as well as across all autosomal genes as a reference value).

| Trait | Variant | | Model | OR (95% CI) or $\beta$ (s.e.) | $P$ |
|---|---|---|---|---|---|
| **(a) 7q22.1** | | | | | |
| T2D | 99kb segment CN | Cont | linear | 0.001 (0.0002) | $2.4 \times 10^{-13}$ |
| T2D | *RASA4* Y731C CN | Cont | linear | 0.004 (0.0004) | $1.3 \times 10^{-25}$ |
| T2D | *RASA4* Y731C CN | Cat | logistic | 1.09 (1.07–1.11) | $2.5 \times 10^{-25}$ |
| T2D | chr7:102579109:G CN | Cont | linear | 0.002 (0.0002) | $1.5 \times 10^{-14}$ |
| Chronotype | 99kb segment CN | Cont | linear | 0.012 (0.0009) | $1.2 \times 10^{-43}$ |
| Chronotype | *RASA4* Y731C CN | Cont | linear | 0.034 (0.002) | $2.6 \times 10^{-72}$ |
| Chronotype | *RASA4* Y731C CN | Cat | linear | 0.034 (0.002) | $3.5 \times 10^{-72}$ |
| Chronotype | chr7:102579109:G CN | Cont | linear | 0.021 (0.001) | $1.2 \times 10^{-73}$ |
| **(b) *CTRB2*** | | | | | |
| T2D | chr16:75204700-75205000 | Cont | linear | – | $1.6 \times 10^{-16}$ |
| T2D | *CTRB2* exon 6 DEL | Cat | logistic | 0.86 (0.82–0.89) | $2.4 \times 10^{-14}$ |
| Pancreatic cancer | chr16:75204700-75205000 | Cont | linear | – | $4.2 \times 10^{-12}$ |
| Pancreatic cancer | *CTRB2* exon 6 DEL | Cat | logistic | 1.46 (1.31–1.62) | $2.5 \times 10^{-12}$ |
| **(c) *FCGR3B*** | | | | | |
| Basophil ct. | *FCGR3B* CN | Cat | linear | 0.06 (0.003) | $1.4 \times 10^{-82}$ |
| Basophil ct. | *FCGR3A* CN | Cat | linear | 0.01 (0.005) | 0.008 |
| COPD | *FCGR3B* CN | Cat | logistic | 0.93 (0.90–0.96) | $7.5 \times 10^{-7}$ |
| RA | *FCGR3B* CN | Cat | logistic | 0.91 (0.88–0.95) | $2.2 \times 10^{-5}$ |
| **(d) *DEFA1/DEFA3*** | | | | | |
| Basophil ct. | 19kb segment CN | Cont | linear | 0.0002 (0.006) | 0.98 |
| Basophil ct. | 5-SNP haplotype CN | Cont | linear | –0.02 (0.001) | $1.0 \times 10^{-75}$ |
| Monocyte ct. | 19kb segment CN | Cont | linear | 0.05 (0.007) | $4.5 \times 10^{-13}$ |
| Monocyte ct. | 5-SNP haplotype CN | Cont | linear | 0.02 (0.001) | $5.9 \times 10^{-72}$ |
| 5-SNP haplotype : **6993547:C/A**; 6993555:G/A; 6993595:C/T; 6993602:C/T; 6993671:G/A | | | | | |

**Supplementary Table 6. Detailed association statistics at highlighted loci involving common CNVs.** (a) 7q22.1 segmental duplication locus associated with T2D and chronotype. Copy numbers of paralogous sequence variants (PSVs) witin the 99kb repeat associated much more strongly with both traits than copy number of the segment. For chronotype, the top-associated variant was the PSV chr7:102579109:G (GRCh38 coordinates for the PSV position within one unit of the 99kb segmental duplication), which slightly beat the *RASA4* Y731C missense PSV. (b) *CTRB2* exon 6 deletion associated with T2D and pancreatic cancer. Association data are shown for a continuous-valued measurement of relative copy number derived from normalized WES read counts at chr16:75204700-75205000 (counting reads with any mapping quality) as well as for hard-called 0/1/2 genotypes (used to obtain interpretable odds ratios). (c) Fc-gamma receptor locus. The association with basophil count was specific to *FCGR3B* copy-number (and not *FCGR3A*), and *FCGR3B* copy-number also associated with rheumatoid arthritis (as previously reported) and COPD. (d) *DEFA1 / DEFA3* segmental duplication locus associated with basophil and monocyte counts. The strongest associations involved copy-number of PSVs within a 5-SNP haplotype in an Alu element within the 19kb repeat unit (Section 9.3); association data are shown for the PSV in bold. Cont, continuous-valued; Cat, categorized into discrete (integer) copy numbers. For linear models, effect size ($\beta$) and standard error (s.e.) are provided in units of standard deviations; for logistic models, odds ratio (OR) and 95% confidence intervals are provided. All models controlled for WES release, assessment center, genotype array, sex, age, age squared, and 20 PCs.

| (a) Quantitative trait | $\beta$ | SE | $P$ | $P$, BOLT-LMM |
|---|---|---|---|---|
| HLS Retic. Count | -0.015 | 0.0029 | $1.3 \times 10^{-7}$ | $1.5 \times 10^{-8}$ |
| Mean Platelet Vol. | 0.013 | 0.0028 | $3.3 \times 10^{-6}$ | $4.5 \times 10^{-9}$ |
| Mean Sphered Cell Vol. | 0.015 | 0.0028 | $2.9 \times 10^{-8}$ | $3.5 \times 10^{-10}$ |
| Basophil Count | 0.016 | 0.0026 | $2.6 \times 10^{-9}$ | $6.6 \times 10^{-11}$ |
| RBC Distr. Width | 0.016 | 0.0028 | $1.6 \times 10^{-8}$ | $4.8 \times 10^{-10}$ |
| White Count | 0.016 | 0.0028 | $4.9 \times 10^{-9}$ | $7.6 \times 10^{-17}$ |
| Eosinophil Count | 0.017 | 0.0028 | $3.3 \times 10^{-9}$ | $5.1 \times 10^{-17}$ |
| Lymphocyte Count | 0.021 | 0.0028 | $3.4 \times 10^{-14}$ | $1 \times 10^{-22}$ |
| HbA1c | 0.027 | 0.0029 | $7.1 \times 10^{-21}$ | $7.6 \times 10^{-30}$ |
| Calcium | 0.034 | 0.003 | $5.6 \times 10^{-31}$ | $1.7 \times 10^{-37}$ |

| (b) GTEx tissue | Genotype counts | $\frac{SIGLEC14}{SIGLEC5}$ TPM | *SIGLEC5* aFC (95% CI) | *SIGLEC5* NES (SE;p) | *SIGLEC14* aFC (95% CI) | *SIGLEC14* NES (SE;p) |
|---|---|---|---|---|---|---|
| Whole_Blood | 434/195/30 | 0.67 | 0.52 (0.45-0.6) | -0.38 (0.03;0) | 0 (0-0.04) | -1.06 (0.02;0) |
| Brain_Cerebellum | 138/63/8 | 1.29 | 1.55 (1.07-3) | 0.1 (0.12;0.43) | 0.21 (0-0.84) | -0.77 (0.1;0) |
| Brain_Cerebellar_Hemisphere | 111/56/8 | 1.55 | 1.53 (1.18-1.97) | 0.19 (0.12;0.1) | 0.11 (0-0.34) | -0.81 (0.08;0) |
| Kidney_Cortex | 48/18/6 | 1.95 | 2.28 (1.49-3.96) | 0.32 (0.15;0.05) | 0 (0-0) | -1.1 (0.11;0) |
| Brain_Nucleus_accumbens_basal_ganglia | 126/66/9 | 1.96 | 1.81 (1.41-2.37) | 0.21 (0.1;0.03) | 0.06 (0-0.27) | -0.79 (0.07;0) |
| Liver | 136/52/15 | 2.01 | 1.28 (0.97-1.65) | 0.12 (0.09;0.19) | 0 (0-0.16) | -0.71 (0.06;0) |
| Brain_Cortex | 129/68/6 | 2.18 | 1.28 (1.02-1.62) | 0.25 (0.11;0.03) | 0 (0-0.02) | -0.99 (0.07;0) |
| Brain_Putamen_basal_ganglia | 113/51/6 | 2.26 | 1.35 (1.08-1.72) | 0.32 (0.12;0.01) | 0 (0-0.08) | -0.84 (0.07;0) |
| Brain_Caudate_basal_ganglia | 122/64/8 | 2.30 | 1.66 (1.22-2.84) | 0.19 (0.1;0.07) | 0.06 (0-0.28) | -0.92 (0.07;0) |
| Brain_Frontal_Cortex_BA9 | 108/55/10 | 2.33 | 3.55 (2.11-8) | 0.16 (0.09;0.09) | 0 (0-0.16) | -0.82 (0.07;0) |
| Thyroid | 365/172/25 | 2.47 | 1.11 (0.96-1.32) | 0.12 (0.06;0.04) | 0 (0-0.06) | -0.86 (0.04;0) |
| Spleen | 144/65/12 | 2.52 | 1.19 (1.07-1.35) | 0.22 (0.06;0) | 0 (0-0.03) | -1.14 (0.04;0) |
| Pituitary | 148/73/12 | 2.56 | 1.56 (1.24-1.92) | 0.2 (0.09;0.02) | 0.2 (0-0.42) | -0.82 (0.06;0) |
| Muscle_Skeletal | 463/199/32 | 2.74 | 1.06 (0.86-1.3) | 0 (0.06;1) | 0.08 (0-0.2) | -0.74 (0.04;0) |
| Heart_Left_Ventricle | 245/115/17 | 2.78 | 1.18 (0.91-1.63) | 0.04 (0.07;0.54) | 0 (0-0.07) | -0.96 (0.05;0) |
| Brain_Anterior_cingulate_cortex_BA24 | 89/52/5 | 2.78 | 1.63 (1-2.93) | 0.06 (0.12;0.61) | 0.22 (0-0.53) | -0.71 (0.08;0) |
| Lung | 325/157/23 | 2.84 | 1.39 (1.19-1.66) | 0.27 (0.05;0) | 0 (0-0.03) | -1.05 (0.04;0) |
| Adipose_Visceral_Omentum | 300/138/23 | 2.88 | 1.19 (1.01-1.47) | 0.19 (0.04;0) | 0 (0-0.05) | -0.87 (0.03;0) |
| Skin_Sun_Exposed_Lower_leg | 381/184/30 | 2.91 | 1.36 (1.17-1.56) | 0.23 (0.05;0) | 0 (0-0.04) | -0.84 (0.04;0) |
| Skin_Not_Sun_Exposed_Suprapubic | 329/151/24 | 2.95 | 1.22 (0.99-1.51) | 0.12 (0.06;0.05) | 0.04 (0-0.16) | -0.81 (0.04;0) |
| Esophagus_Mucosa | 314/152/20 | 2.97 | 1.2 (0.75-1.56) | 0.26 (0.06;0) | 0 (0-0.08) | -0.76 (0.04;0) |
| Prostate | 143/63/11 | 3.18 | 1.66 (1.38-1.98) | 0.38 (0.09;0) | 0 (0-0.09) | -0.91 (0.06;0) |
| Heart_Atrial_Appendage | 232/113/19 | 3.22 | 1.25 (0.98-1.51) | 0.25 (0.06;0) | 0 (0-0.05) | -0.83 (0.04;0) |
| Brain_Hypothalamus | 104/55/10 | 3.26 | 2.15 (1.63-3) | 0.35 (0.11;0) | 0 (0-0.09) | -0.9 (0.07;0) |
| Stomach | 212/91/14 | 3.27 | 1.09 (0.82-1.41) | 0.12 (0.07;0.09) | 0 (0-0.04) | -0.94 (0.04;0) |
| Minor_Salivary_Gland | 84/48/5 | 3.31 | 3.23 (1.78-5.97) | 0.34 (0.09;0) | 0 (0-0.07) | -0.91 (0.08;0) |
| Artery_Tibial | 378/167/27 | 3.38 | 1.63 (1.37-1.92) | 0.36 (0.06;0) | 0 (0-0.06) | -0.88 (0.03;0) |
| Brain_Amygdala | 79/43/5 | 3.42 | 3.45 (2.1-8.42) | 0.33 (0.11;0) | 0 (0-0.15) | -0.9 (0.08;0) |
| Breast_Mammary_Tissue | 247/121/20 | 3.46 | 1.55 (1.32-1.81) | 0.37 (0.07;0) | 0.05 (0-0.16) | -0.83 (0.04;0) |
| Uterus | 82/37/10 | 3.55 | 1.98 (1.38-3.83) | 0.43 (0.12;0) | 0.05 (0-0.21) | -0.85 (0.1;0) |
| Brain_Hippocampus | 101/55/8 | 3.57 | 1.63 (1.18-2.26) | 0.25 (0.1;0.02) | 0 (0-0.11) | -0.95 (0.07;0) |
| Vagina | 93/35/10 | 3.59 | 1.93 (1.35-2.59) | 0.38 (0.12;0) | 0.34 (0.03-0.67) | -0.75 (0.08;0) |
| Small_Intestine_Terminal_Ileum | 108/54/9 | 3.60 | 1.66 (1.38-2.02) | 0.26 (0.06;0) | 0 (0-0.04) | -0.92 (0.05;0) |
| Adipose_Subcutaneous | 376/164/28 | 3.65 | 1.41 (1.23-1.61) | 0.29 (0.06;0) | 0 (0-0.03) | -0.98 (0.03;0) |
| Ovary | 104/51/10 | 3.65 | 1.4 (1.02-1.87) | 0.16 (0.1;0.1) | 0.06 (0-0.3) | -0.7 (0.06;0) |
| Colon_Transverse | 225/116/17 | 3.69 | 1.33 (1.14-1.56) | 0.22 (0.06;0) | 0 (0-0.05) | -0.88 (0.04;0) |
| Pancreas | 190/91/17 | 3.79 | 1.56 (1.21-1.96) | 0.32 (0.07;0) | 0 (0-0.01) | -0.96 (0.05;0) |
| Artery_Coronary | 131/63/13 | 3.81 | 1.76 (1.48-2.14) | 0.33 (0.08;0) | 0 (0-0.1) | -0.8 (0.05;0) |
| Esophagus_Gastroesophageal_Junction | 203/102/18 | 3.92 | 1.3 (1.08-1.55) | 0.26 (0.07;0) | 0.02 (0-0.13) | -0.86 (0.04;0) |
| Artery_Aorta | 246/112/21 | 3.92 | 1.4 (1.09-1.77) | 0.27 (0.05;0) | 0 (0-0.07) | -0.83 (0.04;0) |
| Esophagus_Muscularis | 293/142/20 | 4.02 | 1.49 (1.24-1.75) | 0.32 (0.06;0) | 0.01 (0-0.11) | -0.91 (0.03;0) |
| Testis | 211/92/10 | 4.05 | 1.58 (1.45-1.74) | 0.72 (0.09;0) | 0.02 (0-0.08) | -1.18 (0.05;0) |
| Colon_Sigmoid | 206/91/12 | 4.22 | 1.76 (1.37-2.25) | 0.38 (0.09;0) | 0.04 (0-0.22) | -0.87 (0.05;0) |
| Brain_Spinal_cord_cervical_c-1 | 79/39/7 | 4.36 | 4.12 (2.83-7.9) | 0.52 (0.09;0) | 0 (0-0.08) | -0.89 (0.07;0) |
| Nerve_Tibial | 347/150/26 | 4.66 | 1.77 (1.59-1.97) | 0.5 (0.06;0) | 0.05 (0-0.13) | -0.96 (0.03;0) |
| Brain_Substantia_nigra | 68/40/5 | 4.66 | 3.78 (2.75-5.67) | 0.72 (0.11;0) | 0.11 (0-0.36) | -0.83 (0.08;0) |
| Cells_EBV-transformed_lymphocytes | 90/50/4 | 7.19 | 9.59 (6.35-22.73) | 1.21 (0.1;0) | 0 (0-0.07) | -0.93 (0.1;0) |
| Adrenal_Gland | 148/67/15 | 7.29 | 2.3 (1.92-2.76) | 0.58 (0.09;0) | 0 (0-0.01) | -1.05 (0.05;0) |

**Supplementary Table 7. Detailed association statistics for *SIGLEC14–SIGLEC5* gene fusion.** (a) Quantitative trait associations for the *SIGLEC14–SIGLEC5* fusion. Effect size, standard error and $P$-value are from linear regression on 0/1/2 genotypes (excluding individuals with reciprocal duplications from analysis) controlling for WES release, assessment center, genotype array, sex, age, age squared, and 20 PCs. The linear mixed model $P$-value from BOLT-LMM for the continuous-valued measurement of relative copy number derived from normalized WES read counts at chr19:51630200-51646900 (counting reads with any mapping quality) is also provided. (b) Allelic fold change (aFC) and normalized effect size (NES) of the *SIGLEC14–SIGLEC5* fusion on *SIGLEC5* and *SIGLEC14* expression in GTEx tissues. For each tissue, the median ratio of *SIGLEC14* TPM and *SIGLEC5* TPM (among individuals with no gene fusion or *SIGLEC14* duplication)—measuring the relative efficiency of the *SIGLEC14* vs. *SIGLEC5* promoters—is also provided, along with counts of hom-REF/het-fusion/hom-fusion genotypes per tissue.
